# Large uncertainty in individual PRS estimation impacts PRS-based risk stratification

**DOI:** 10.1101/2020.11.30.403188

**Authors:** Yi Ding, Kangcheng Hou, Kathryn S. Burch, Sandra Lapinska, Florian Privé, Bjarni Vilhjálmsson, Sriram Sankararaman, Bogdan Pasaniuc

**Affiliations:** Bioinformatics Interdepartmental Program, UCLA, Los Angeles, CA 90095; Department of Statistics and Data Science, Cornell University, Ithaca, NY 14853; Department of Economics and Business Economics, National Centre for Register-Based Research, Aarhus University, Aarhus, Denmark; Department of Computer Science, UCLA, Los Angeles, CA 90095; Department of Computational Medicine, David Geffen School of Medicine at UCLA, Los Angeles, CA 90095; Department of Human Genetics, David Geffen School of Medicine at UCLA, Los Angeles, CA 90095; Department of Pathology and Laboratory Medicine, David Geffen School of Medicine at UCLA, Los Angeles, CA 90095

## Abstract

Large-scale genome-wide association studies have enabled polygenic risk scores (PRS), which estimate the genetic value of an individual for a given trait. Since PRS accuracy is typically assessed using cohort-level metrics (e.g., R^2^), uncertainty in PRS estimates at individual level remains underexplored. Here we show that Bayesian PRS methods can estimate the variance of an individual’s PRS and can yield well-calibrated credible intervals for the genetic value of a single individual. For real traits in the UK Biobank (N=291,273 unrelated “white British”) we observe large variance in individual PRS estimates which impacts interpretation of PRS-based stratification; for example, averaging across 13 traits, only 0.8% (s.d. 1.6%) of individuals with PRS point estimates in the top decile have their entire 95% credible intervals fully contained in the top decile. We provide an analytical estimator for individual PRS variance—a function of SNP-heritability, number of causal SNPs, and sample size—and observe high concordance with individual variances estimated via posterior sampling. Finally as an example of the utility of individual PRS uncertainties, we explore a probabilistic approach to PRS-based stratification that estimates the probability of an individual’s genetic value to be above a prespecified threshold. Our results showcase the importance of incorporating uncertainty in individual PRS estimates into subsequent analyses.

## Introduction

Polygenic risk scores (PRS) have emerged as the main approach for predicting the genetic component of an individual’s phenotype and/or common-disease risk (i.e. genetic value, GV) from large-scale genome-wide association studies (GWAS). Several studies have demonstrated the utility of PRS as estimators of genetic values in genomic research and, when combined with non-genetic risk factors (e.g., age, diet, etc), in clinical decision-making^1–3^—for example, in stratifying patients^4^, delivering personalized treatment^5^, predicting disease risk^6^, forecasting disease trajectories^7,8^, and studying shared etiology among traits^9,10^. Increasingly large GWAS sample sizes have improved the predictive value of PRS for several complex traits and diseases^11,12^ including breast cancer^6,13^, prostate cancer^14^, lung cancer^15^, coronary artery disease^16^, obesity^7^, type 1 diabetes^17^, type 2 diabetes^18^, and Alzheimer’s disease^19^, thus paving the way for PRS-informed precision medicine.

Under a linear additive genetic model, an individual’s genetic value (GV; the estimand of interest for PRS) is the sum of the individual’s dosage genotypes at causal variants (encoded as the number of copies of the effect allele) weighted by the causal allelic effect sizes (expected change in phenotype per copy of the effect allele). In practice, the true causal variants and their effect sizes are unknown and must be inferred from GWAS data. Existing PRS methods generally fall into one of three categories based on their inference procedure: (1) pruning/clumping and thresholding (P+T) approaches, which account for linkage disequilibrium (LD) by pruning/clumping variants at a given LD and/or significance threshold and weight the remaining variants by their marginal association statistics^20,21^; (2) methods that account for LD through regularization of effect sizes, including lassosum^22^ and BLUP prediction^23,24^; and (3) Bayesian approaches that explicitly model causal effects and LD to infer the posterior distribution of causal effect sizes^25–27^.

Both the bias and variability of a PRS estimator are critical to assessing its practical utility. Given that most PRS methods select variants (predictors) and estimate their effect sizes, there are two main sources of uncertainty: (1) uncertainty about which variants are causal (i.e. have non-zero effects) and (2) statistical noise in the causal effect estimates due to the finite sample size of GWAS training data. The impact of sample size and LD on causal variant identification has been thoroughly investigated in the statistical fine-mapping literature^28,29^, with uncertainty increasing as the strength of LD in a region increases and as the sample size of the GWAS training data decreases. As a toy example, consider a region with two variants with same marginal GWAS statistics that are in near-perfect LD: without additional information, it is impossible to determine whether one or both of the variants are causal given finite sample size and small effect sizes^28,29^. This uncertainty about which variant is causal propagates into uncertainty in the weights used for PRS, leading to different estimates of genetic value in a target individual. Evaluating how this uncertainty propagates to individual PRS estimation may improve subsequent analyses such as PRS-based risk stratification.

Unfortunately, studies that have applied PRS and/or examined PRS accuracy have largely ignored uncertainty in PRS estimates at the individual level^1^, focusing instead on cohort-level metrics of accuracy such as R^2^. Therefore, the degree to which uncertainty in causal variant identification impacts individual PRS estimation and subsequent analyses (e.g., stratification) remains unclear. In contrast, in livestock breeding programs, prediction error variance (PEV) of estimated breeding values has been used for decades to evaluate the precision of individual estimated breeding values and to generate other genetic evaluation statistics^30–32^. PEV can be directly computed by inverting the coefficient matrix of mixed model equations^30,33^ or, if inversion is computationally prohibitive, approximated^34–39^. The uncertainty in other biomarkers and non-genetic risk factors have also been well-studied^40^. For example, smoothing methods and error-correction methods are performed before biomarkers and non-genetic risk factors are included in the predictive model^41,42^.

Motivated by potential clinical applications of PRS in personalized medicine, where one of the main goals is to estimate risk of a given individual, we focus on evaluating uncertainty in PRS estimates at the level of a single target individual. Our goal is to quantify the statistical noise in individual PRS estimates 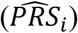 conditional on data used to train the PRS. We assess two metrics of individual PRS uncertainty: (1) the standard deviation of the PRS estimate for individual *i*, denoted 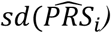; and (2) the *ρ*-level credible interval for the genetic value of individual *i*, defined as the interval that contains the genetic value of individual *i* (GV_i_) with *ρ* (e.g., 95%) probability, denoted (*ρ* GV_i_-CI). We extend the Bayesian framework of LDpred2^24^, a widely used method for PRS estimation, to sample from the posterior distribution of Gv_*i*_ to estimate 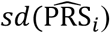 and *ρ* GV_i_-CI for different values of *ρ*. First, we introduce an analytical form for the expectation across individuals of 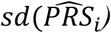 as function of heritability, number of causals and training data sample size and show that the analytical form is accurate in simulations and real data. Second, we use simulations starting from real genotypes in the UK Biobank (N=291,273 individuals, M=459,792 SNPs, unrelated “white British”) to show that *ρ* GV_i_-CI is well-calibrated when the target sample matches the training data and that 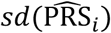 increases as polygenicity (number of causal variants) increases and as heritability and GWAS sample size decrease^43^. Analyzing 13 real traits in the UK Biobank, we observe large uncertainties in individual PRS estimates that greatly impact the interpretability of PRS-based ranking of individuals. For example, on average across traits, only 0.2% (s.d. 0.6%) of individuals with PRS point estimates in the top 1% also have corresponding 95% GV_i_-CI fully contained in the top 1%. Individuals with PRS point estimates at the 90^th^ percentile in a testing sample can be ranked anywhere between the 34^th^ and 99^th^ percentiles in the same testing sample after their 95% credible intervals are taken into account. Finally, we explore a probabilistic approach to incorporating PRS uncertainty in PRS-based stratification and demonstrate how such approaches can enable principled risk stratification under different cost scenarios.

## Results

### Sources of uncertainty in individual PRS estimation

Under a standard linear model relating genotype to phenotype (Methods), the estimand of interest for PRS is the genetic value of an individual *i*, defined as 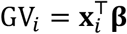, where **x**_*i*_ is an *M* × 1 vector of observed genotypes and **β** is the corresponding *M* × 1 vector of unknown causal effect sizes^44^ (Methods). Different PRS methods vary in how they estimate causal effects 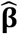 to construct the estimator 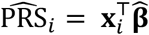. Inferential variance in 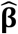 propagates into the variance of 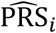. In this work, we focus on quantifying the inferential uncertainty in 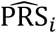 and assessing its impact on PRS-based stratification.

To illustrate the impact of statistical noise in 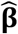 on 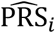, consider a toy example of a trait for which the observed marginal GWAS effects at three SNPs are equal (Figure 1). The trait was simulated assuming SNP1 and SNP2 are causal with the same effect whereas SNP3 is not causal but tags SNP2 with high LD (0.9). The *expected* marginal effect is higher at SNP2 than at SNP3, thus implying that GWAS with infinite sample size would correctly identify the true causal variants and their effects. However, finite GWAS sample sizes induce statistical noise in the *observed* marginal effects; for example, the marginal effect at SNP3 (tag SNP) is higher than at SNP1 (true causal SNP) in 12% to 30% of GWASs simulated with sample size N=100,000 under the LD structure of Figure 1 (Supplementary Figure 1). Thus, the key challenge is that, given only GWAS marginal effects and LD, there is more than one plausible causal effect-size configuration. In Figure 1, the observed marginal effects (the same at all three SNPs) could be driven by SNPs (1 and 2) or (1 and 3) or (1, 2, and 3); in fact, (1 and 2) and (1 and 3) are equally probable in absence of other information. In such situations, one can generate different PRS estimates for a given individual from the same training data. For example, P+T PRS methods and lassosum, which assume sparsity, would likely select either SNPs (1 and 2) or (1 and 3), while BLUP or Bayesian approaches would likely take an average over the possible causal configurations, splitting the causal effect of SNP2 between SNPs (2 and 3). Thus, in such cases, an individual with the genotype **x**_*i*_ = (0,1,0)^T^ can be classified as being above or below a prespecified threshold, depending on the approach/assumptions used to estimate causal effects.

**Figure 1.**
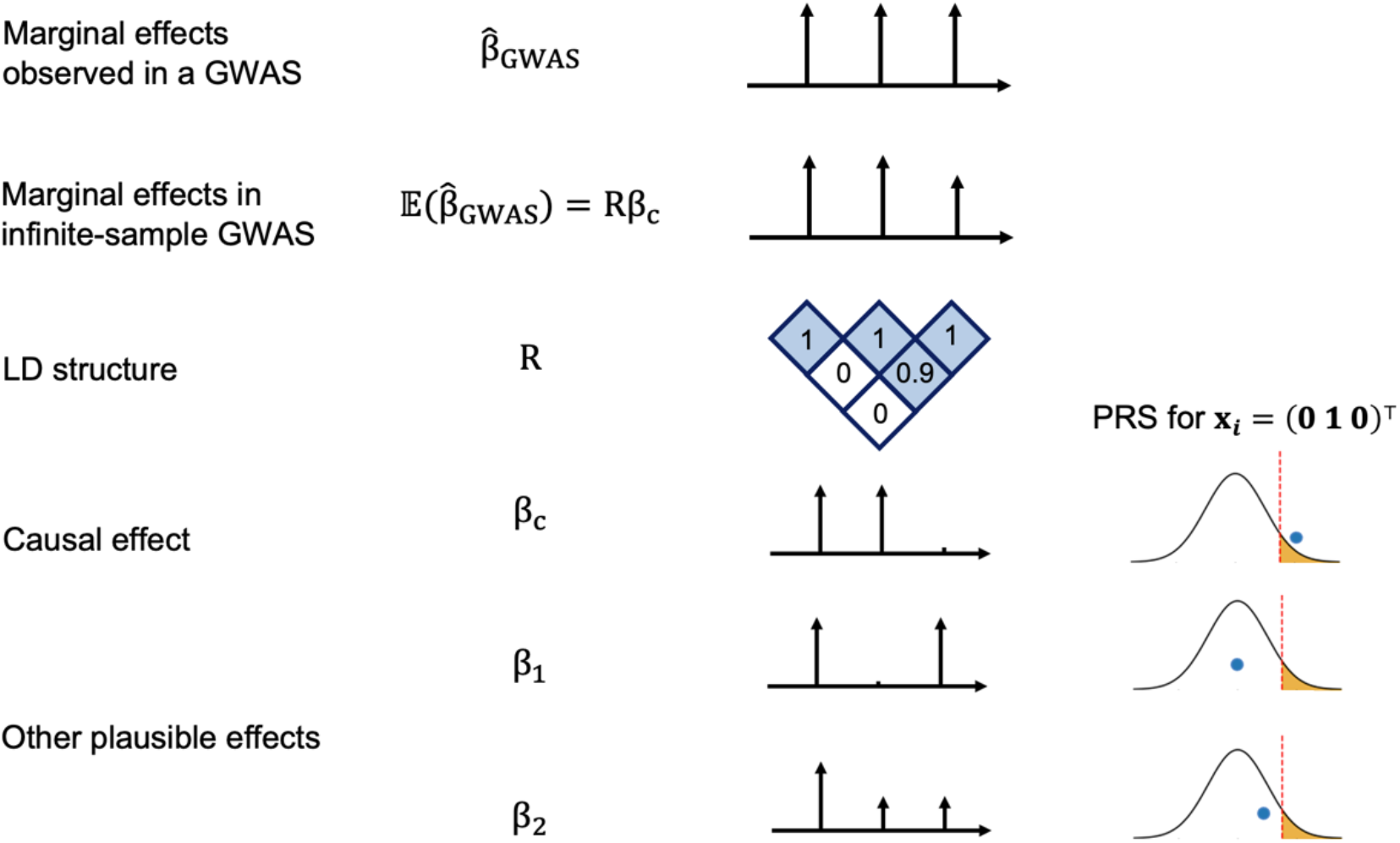
LD and finite GWAS sample size introduce uncertainty into PRS estimation. We simulated a GWAS of **N** individuals across 3 SNPs with LD structure **R** (SNP2 and SNP3 are in LD of 0.9 whereas SNP1 is uncorrelated to other SNPs) where SNP1 and SNP2 are causal with the same effect size **β**_**c**_ = (0.016, 0.016, 0) such that the variance explained by this region is var(**x**^T^**β**_**c**_) = 0.5/1000 corresponding to a trait with total heritability of 0.5 uniformly distributed across 1,000 causal regions. The marginal effects observed in a GWAS, 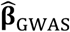, have an expectation of **Rβ**_**c**_ and variance-covariance 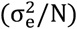**R**, thus showcasing the statistical noise introduced by finite sample size of GWAS (N); for example, the probability of the marginal GWAS effect at tag SNP3 to exceed the marginal effect of true causal SNP2, although decreases with N, remains considerably high for realistic sample and effect sizes (12% at N=100,000 for a trait with h2=0.5 split across 1,000 causal regions, see Supplementary Figure 1). We consider one such observation for the effects observed in a GWAS: 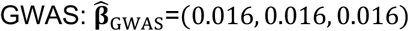. Given such observation, in addition to the true causal effects (**β**_**c**_), other causal configurations are probable **β**_**1**_ **=** (0.016, 0, 0.016) or **β**_**2**_ **=** (0.016, 0.008, 0.008). An individual with genotype **x**_**i**_ = (**0 1 0**)^T^ will attain different PRS estimates under these different causal configurations. Most importantly, in the absence of other prior information, **β**_**1**_ and **β**_**c**_ are equally probable given the data thus leading to different PRS estimates for individual **x**_**i**_ = (**0 1 0**)^T^.

We explore inferential uncertainty in 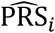 through two synergistic approaches. First, we provide a closed-form approximation for the expected 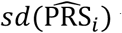 under simplifying assumptions. Second, we sample from the posterior distribution of the causal effects under the framework of LDPred2 to estimate 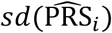 and compute credible intervals for GV_*i*_ at prespecified confidence levels (e.g., *ρ* = 95%) (Figure 2). As an example of the utility of such measures of uncertainty, we explore a probabilistic approach to PRS-based risk stratification that estimates the probability that GV_*i*_ is above a given threshold *t* (Figure 2) and demonstrate how this probability can be used in conjunction with situation-specific cost functions to optimize risk stratification decisions.

**Figure 2.**
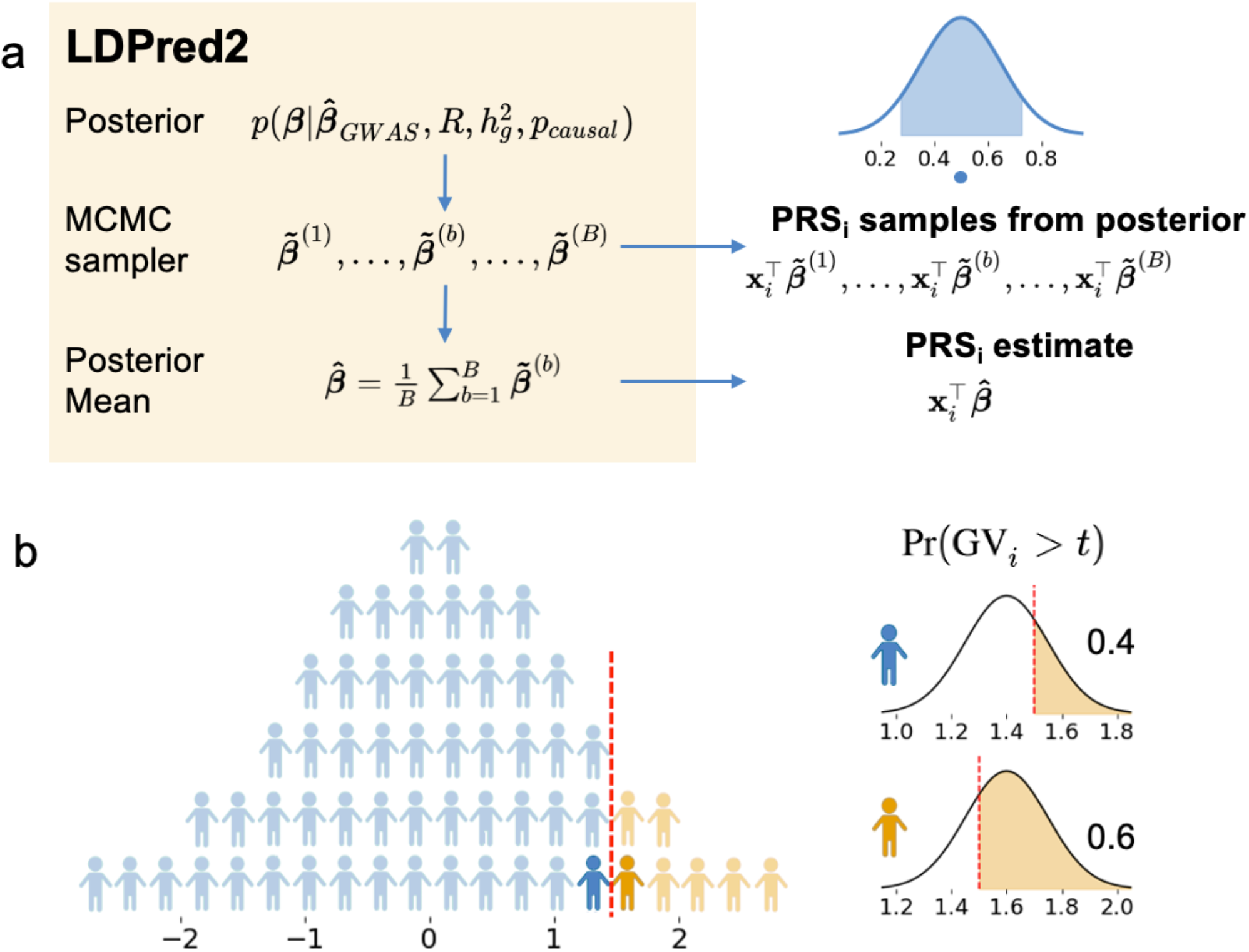
Framework for extracting uncertainty from Bayesian methods for probabilistic individual stratification. (a) Procedure to obtain uncertainty from LDpred2. LDpred2 uses MCMC to sample from the posterior causal effect distribution given GWAS marginal effects, LD, and a prior on the causal effects. It outputs the posterior mean of the causal effects which is used to estimate the posterior mean genetic value (the PRS point estimate). Our framework samples from the posterior of the causal effects to approximate the posterior distribution of genetic value. The density plot represents the posterior distribution of GV for an individual. The shaded area represents a ρ-level credible interval. The dot represents the posterior mean. (b) Probabilistic risk stratification framework. Given a threshold *t*, instead of dividing individuals into above-threshold 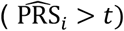 and below-threshold 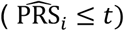 groups dichotomously (left), probabilistic risk stratification assigns each individual a probability of being above-threshold Pr(GV_*i*_ > *t*) (right).

### Analytical derivation of individual PRS uncertainty

We focus on evaluating PRS uncertainty within a general Bayesian framework, where the posterior mean of the genetic effects conditional on a given GWAS, 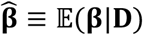, is used to estimate the genetic value of a given individual, 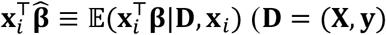 with access to individual data or 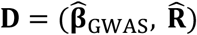 with access to marginal association statistics and LD, see Methods). We define PRS uncertainty for individual *i* as the posterior variance of their genetic value, 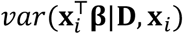. This quantity is an approximation to prediction error variance (PEV) of estimated breeding values (EBV) in livestock genetics^32,34^. EBV is analogous to genetic value in human genetics; derivations relating PRS uncertainty to PEV of EBV can be found in Methods.

Assuming that every SNP has a nonzero causal effect drawn *i*.*i*.*d*. from 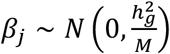, one can derive a closed-form approximation to the expectation across individuals of the posterior variance of genetic value (Methods). Given a GWAS discovery dataset of *N* unrelated individuals drawn from a given population, the expected PRS uncertainty for a test individual *i* randomly drawn from the same population is

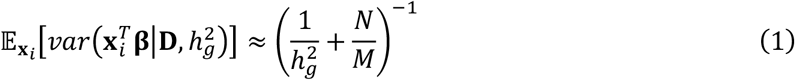

Under an infinitesimal model, the analytical form is an approximately unbiased estimator of the expected posterior variance, even in the presence of LD (Figure 3a). Under non-infinitesimal models, the analytical form underestimates the expected posterior variance, albeit by a relatively small amount (Supplementary Figure 2). Notably, across 13 real phenotypes in the UK Biobank, the analytical form provides relatively accurate estimates of the empirical average 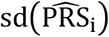 computed from LDpred2 posterior sampling (*R*^2^ = 0.79 across traits, Figure 3b). Thus, the analytical form captures the interplay among SNP-heritability, sample size, and number of causal variants and provides a useful approximation to individual PRS uncertainty when posterior samples are unavailable.

**Figure 3.**
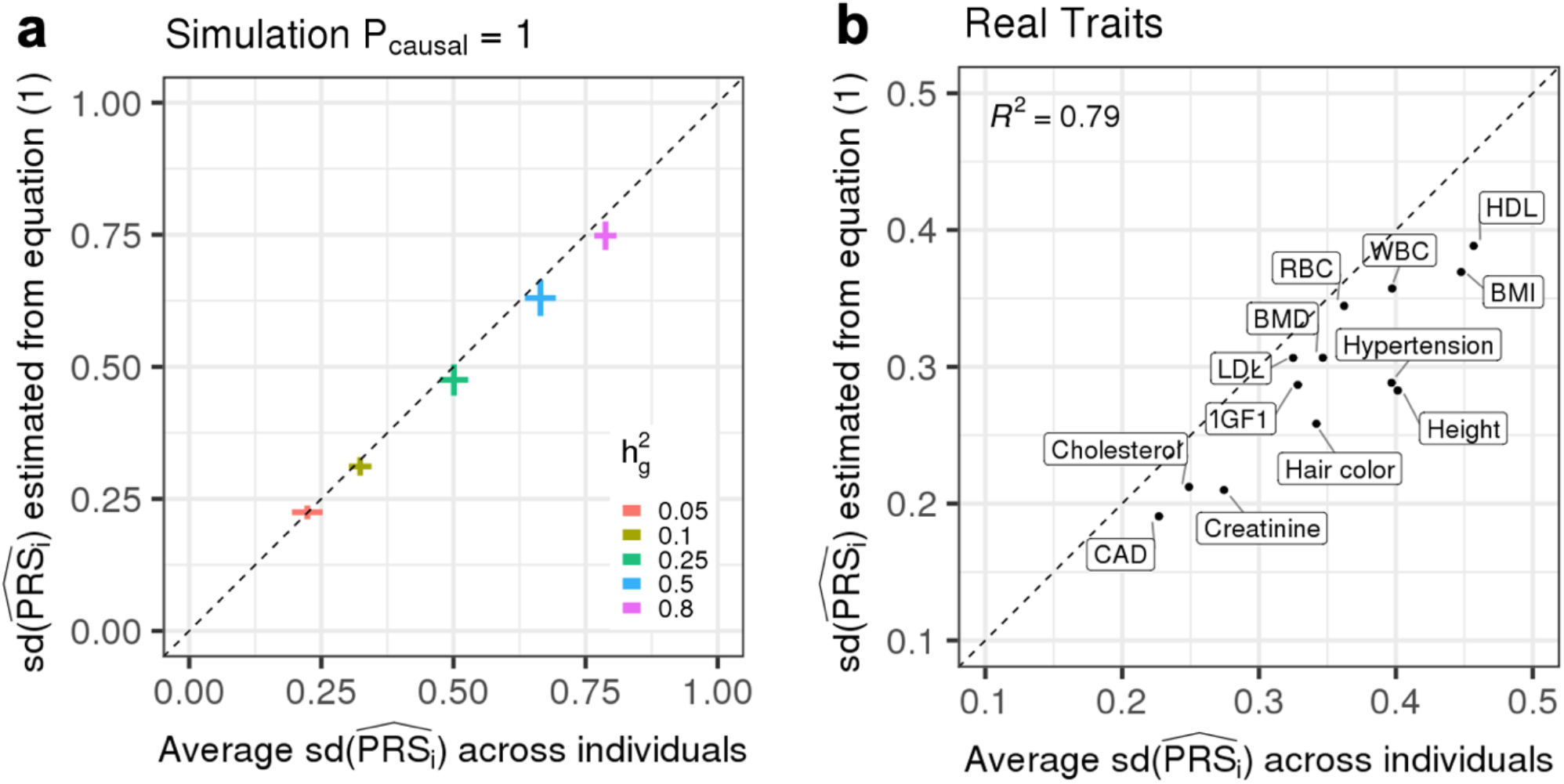
Expected 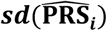 estimated as a function of heritability, polygenicity and training GWAS sample size is highly correlated with average 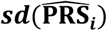 across testing individuals. (a) The analytical form provides approximately unbiased estimates of expected 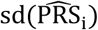 in simulations when p_causal_ = 1. The x-axis is the average 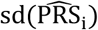 in testing individuals. The y-axis is the expected 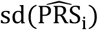 computed from Equation (1). Each dot is an average of 10 simulation replicates for each 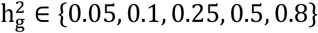. The horizontal whiskers represent ± 1.96 standard deviations of average 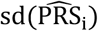 across 10 simulation replicates. The vertical whiskers represent ±1.96 standard deviations of expected 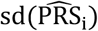 across 10 simulation replicates. (b) The analytical estimator of expected 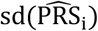 is highly correlated with estimates obtained via posterior sampling for real traits. The x-axis is the average 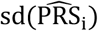 in testing individuals. The y-axis is the expected 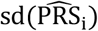 computed from Equation (1), where M is replaced with the estimated number of causal variants and heritability is replaced with estimated SNP-heritability.

### Factors impacting individual PRS uncertainty in simulations

Next, we quantified the degree to which different parameters contribute to uncertainty in individual PRS estimates in simulations starting from real genotypes of unrelated “white British” individuals in the UK Biobank (N=291,273 individuals and M=459,792 SNPs). To avoid overfitting, we partitioned the individuals into disjoint training, validation and testing groups (N_train_=250,000, N_validation_=20,000, N_test_=21,273). Training samples were used to estimate PRS weights; validation samples were used to estimate hyperparameters (e.g., heritability and polygenicity) for LDpred2; and testing samples were used to evaluate accuracy (Supplementary Figure 3) and uncertainty (Methods).

First, we assess the calibration of the *ρ* -level credible intervals for GV_*i*_ estimated by LDpred2. We compared the empirical coverage of the *ρ*-level credible intervals (proportion of individuals in a single simulation replicate whose *ρ* GV_i_-CI overlaps their true GV_*i*_) to the expected coverage (*ρ*) across a range of values of *ρ*. We find that, overall, the *ρ* GV_i_-CI are well-calibrated, albeit slightly mis-calibrated in high-heritability, low-polygenicity simulations (Figure 4a and Supplementary Figure 4). For example, across 10 simulation replicates where 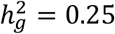 and *p* _causal_= 1%, the 90% GV_i_-CIs have an average empirical coverage of 0.92 (s.e.m. 0.005) (Figure 4a). The *ρ* GV_i_-CIs estimated by LDpred2 are also robust to training cohort sample size (Supplementary Figure 5). Since individuals with large PRS estimates might have larger number of effect alleles and therefore accumulate more inferential variance, we investigate whether individual PRS uncertainty varies with respect to their true genetic value and find no significant correlation between an individual’s 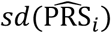 and their true genetic value (Figure 4b).

**Figure 4.**
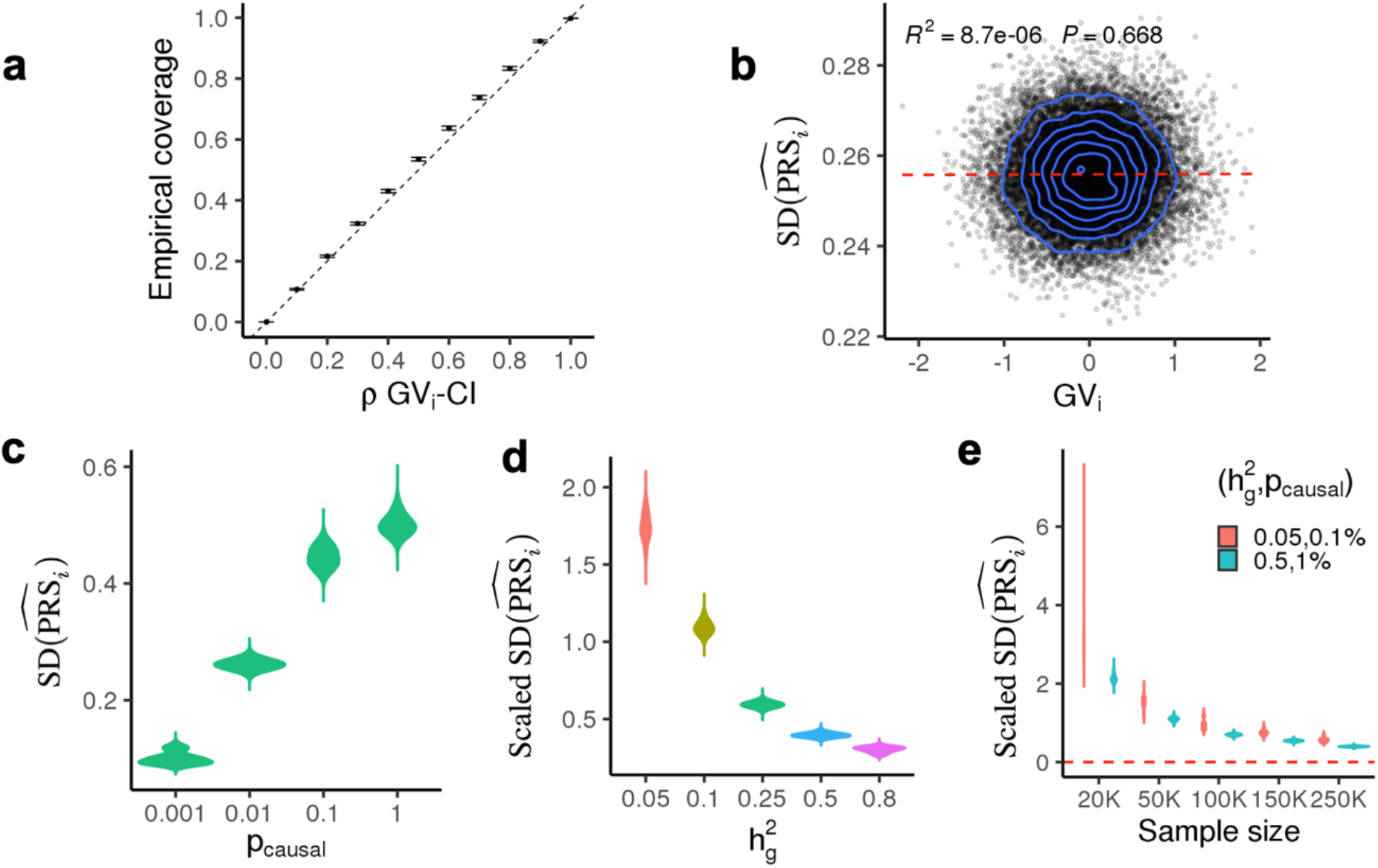
Genetic architecture (polygenicity (*p*_causal_), SNP-heritability 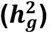, and GWAS sample sizes) impacts uncertainty in PRS estimates in simulations. (a) Individual credible intervals are well-calibrated 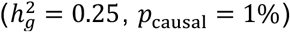. Empirical coverage is calculated as the proportion of individuals in a single simulation whose *ρ*-level credible intervals contain their true genetic risk. The error bars represent 1.96 standard errors of the mean calculated from 10 simulations. (b) Correlation between uncertainty and true genetic value 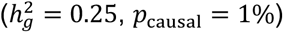. Each dot represents an individual. The x-axis is the true genetic value; the y-axis is standard deviation of the individual PRS estimate 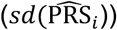. (c) Distribution of individual PRS uncertainty estimates with respect to polygenicity 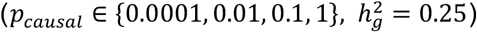. Each violin plot represents 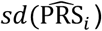 for 21,273 testing individuals across 10 simulations. (d) Distribution of individual PRS uncertainty estimates with respect to heritability 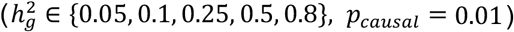. Each violin plot represents scaled 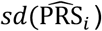 for 21,273 testing individuals across 10 simulation replicates. Since larger heritability yields larger genetic values in our simulations, we plot 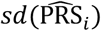 divided by the standard deviation of PRS point estimates in the testing group to enable comparison of uncertainty across different heritability values (Methods). (e) Distribution of individual uncertainty estimates with respect to training GWAS sample size. Each violin plot represents scaled 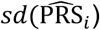 of individual PRS for 21,273 testing individuals across 10 simulation replicates.

We next assessed the impact of trait-specific genetic architecture parameters (heritability and polygenicity) on individual PRS uncertainty, defined as the posterior standard deviation of genetic value. First, we fixed heritability and varied polygenicity and found that 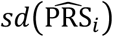 increases from 0.10 to 0.50 when the proportion of causal variants increases from 0.1% to 100% (Figure 4c, Supplementary Figure 6). Second, we varied the heritability while keeping polygenicity constant. Since different heritabilities and sample sizes lead to different variances explained by the PRS in the test sample, we scale the individual standard deviation 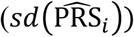 by the standard deviation of PRS point estimates across all tested individuals; we refer to this quantity as “scaled SD” (Methods). We find that the scaled SD decreases with heritability and sample size (Figure 4d, Supplementary Figure 7). For example, when 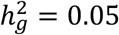 and *p*_causal_ = 0.1%, a 5-fold increase in training data sample size (50K to 250K) reduces scaled SD by 3-fold (from 1.50 to 0.56); when 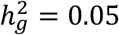 and *p*_causal_ = 1%, the same increase in training data sample size reduces the scaled SD by 4-fold (from 1.10 to 0.39). While the two simulation settings (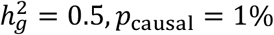 versus 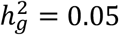 *p*_causal_ = 0.1%) yield the same expected variance per causal variant under our simulation framework (i.e. 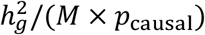, see Methods), we observe lower uncertainty across all sample sizes for 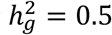 and *p*_causal_ = 1%, further emphasizing the impact of trait-specific genetic architecture on individual PRS uncertainty.

Next, we investigated the impact of different types of model misspecification on credible interval calibration and PRS uncertainty in simulations based on a set of 124,080 SNPs (the union of 36,987 UK Biobank (UKBB) array SNPs and 93,767 HapMap3 SNPs) on chromosome 2. First, we assessed the impact of imperfect tagging of causal variants by simulating phenotypes from the set of HapMap3 + UKBB SNPs 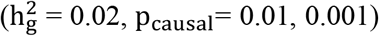 and training the PRS on (i) 124,080 SNPs (HapMap3 + UKBB) and (ii) 36,987 SNPs (UKBB only). The “HapMap3 + UKBB” model contains all causal SNPs whereas the “UKBB only” model excludes ∼70% of the causal SNPs, thus representing imperfect tagging of causal effects. As expected, the empirical coverage of the credible intervals is biased downward across a range of values of *ρ* when only the UKBB SNPs are used to train the model (Supplementary Figure 8). This downward bias is less pronounced when polygenicity is higher (e.g., p_causal_= 0.01 vs 0.001) since the UKBB SNPs tag a larger proportion of heritability due to the increased causal SNP density. Second, to assess whether the coexistence of large and small causal effects impacts PRS uncertainty, we compared three simulation scenarios: (I) large effects only 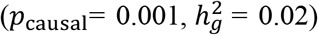, (II) small effects only 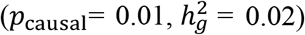, and (III) a “mixture of normal” model (*p*_causal_= 0.0055, 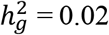 in total) composed of large effects (*p*_causal_= 0.0005, 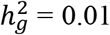) and small effects (*p*_causal_ = 0.005, 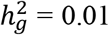). We find that the presence of a large number of small effects increases the uncertainty in individual PRS estimates. For example, the average 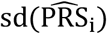 among the 21,273 test individuals is 0.050, 0.087, and 0.11 for simulations I, III and II, respectively (Supplementary Figure 9). In simulation III, both PRS uncertainty and accuracy (squared Pearson correlation between GV and 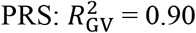, 0.51, 0.68 for I, II, III) are approximate averages of simulations I and II. Despite the LDpred2 model being mis-specified in the mixture of normal simulation, the genetic value credible intervals remain well-calibrated (Supplementary Figure 9). Third, we compared PRS obtained using external reference LD (a subsample of either 1,000 (1K) or 2,000 (2K) individuals held out from the UKBB training data) to those obtained using in-sample LD (all 250,000 individuals in the training data) and found similar degrees of PRS uncertainty and credible interval calibration (Supplementary Figure 10).

### Individual PRS uncertainty in real data in the UK Biobank

We investigate individual PRS uncertainty across 13 traits in the UK Biobank: hair color, height, body mass index (BMI), bone mass density in the heel (BMD), high-density lipoprotein (HDL), low-density lipoprotein (LDL), cholesterol, igf1, creatinine, red blood cell count (RBC), white blood cell count (WBC), hypertension and self-reported cardiovascular disease (CVD). First we focus on PRS-based risk stratification. Since most traits analyzed here are not disease traits, we use “above-threshold” and “below-threshold” when referring to the results of risk stratification. We classify test individuals as above-threshold if their PRS point estimate (the posterior mean of their genetic value) exceeds a prespecified threshold *t* (i.e.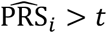), where *t* is set to the 90^th^ PRS percentile obtained from the test-group individuals (Methods). We note that this threshold was chosen arbitrarily to provide an example of how one can compute and interpret PRS uncertainty; in practice, choosing a threshold requires careful consideration of various trait-specific factors such as prevalence and the intended clinical application^1^. We then partition the above-threshold individuals into two categories: individuals whose 95% GV_i_-CI are fully above the threshold *t* (“certain above-threshold”) and individuals whose 95% GV_i_-CI contain *t* (“uncertain above-threshold”). Similarly, we classify individuals as below-threshold if their PRS point estimate lies below a prespecified threshold 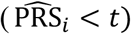 and we partition these individuals into “certain below-threshold” and “uncertain below-threshold” based on their 95% GV_i_-CI (Figure 5a). At *t* = 90^th^ percentile and *ρ* = 95%, only 1.8% (s.d. 2.4%) of above-threshold individuals (averaged across traits) are deemed certain above-threshold individuals; the remaining above-threshold individuals have *ρ*-level credible intervals that overlap *t* (Figure 5b, Table 1). On the other hand, 33.7% (s.d. 15.3%) of below-threshold individuals have *ρ*-level credible intervals that do not overlap *t* (Figure 5b, Table 1). Consistent with simulations, we find that uncertainty is higher for traits that are more polygenic^45^ (Table 1) with the average standard deviation of 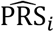 ranging between 0.2 to 0.41 across the studied traits (Table S1). We assessed whether the standard practice of quantile normalization of phenotypes impacts PRS and verify that for phenotypes with mildly skewed distributions, GWAS marginal association statistics and PRS uncertainty are largely consistent with or without quantile normalization (Supplementary Figures 11 and 12).

**Figure 5.**
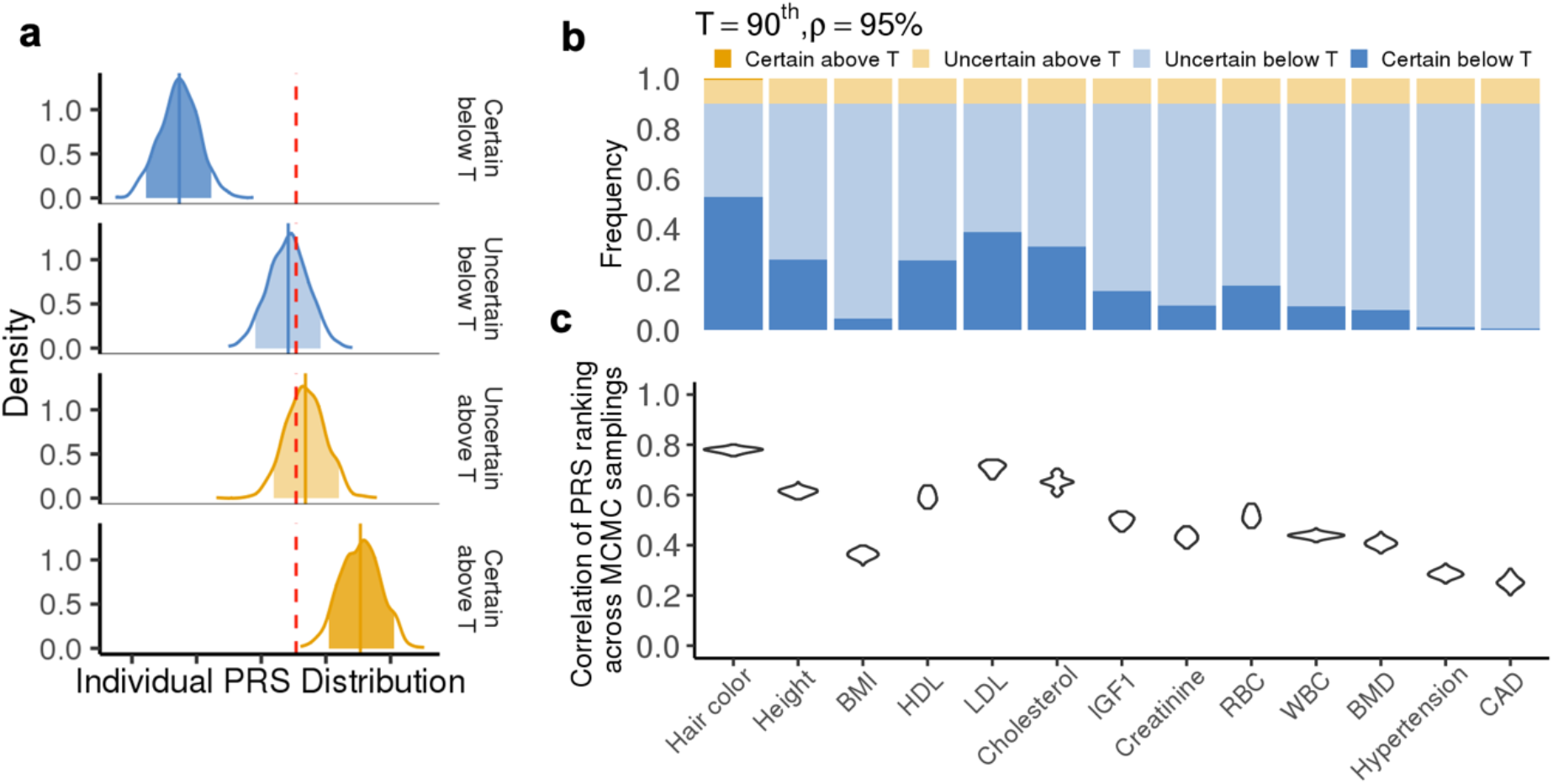
Uncertainty in real data and its influence on genetic risk stratification. (a) Example of posterior PRS distributions for individuals with certain below-threshold (dark blue), uncertain below-threshold (light blue), uncertain above-threshold (light yellow), and certain above-threshold (dark yellow) classifications for HDL. Each density plot is a smoothed posterior PRS distribution of an individual randomly chosen from that category. The solid vertical lines are posterior means. The shaded areas are 95% credible intervals. The red dotted line is the classification threshold. (b) Distribution of classification categories across 11 traits (*t*=90%, *ρ*=95%). Each bar plot represents the frequency of testing individuals who fall into each of the four classification categories for one trait. The frequency is averaged across five random partitions of the whole dataset. (c) Correlation of PRS rankings of test individuals obtained from two MCMC samplings from the posterior of the causal effects. For each trait, we draw two samples from the posterior of the causal effects, rank all individuals in the test data twice based on their PRS from each sample, and compute the correlation between the two rankings across individuals. Each violin plot contains 5,000 points (1,000 pairs of MCMC samples and five random partitions).

**Table 1.** PRS-based individual stratification uncertainty across 11 complex traits in UK Biobank. We quantified PRS-based stratification uncertainty in testing individuals for eleven complex traits at two stratification thresholds (t = 90^th^ and t = 99^th^ percentiles). The numbers of certain versus uncertain classifications are determined from the 95% credible intervals (*ρ* = 95%). For each trait, we report averages (and standard deviations) from five random partitions of the whole dataset.

| Trait | PRS < t (“Below threshold”) |  | PRS > t (“Above threshold”) |  |
| --- | --- | --- | --- | --- |
|  | # *Certain* | # *Certain*/<br>(#*Certain* + #<br>*Uncertain*) | # *Certain* | # *Certain*/<br>(#*Certain* + #<br>*Uncertain*) |
| **t = 90<sup>th</sup>** |  |  |  |  |
| Hair color | 11205.0 (287.0) | 58.5 (1.5)% | 131.4<br>(18.6) | 6.2 (0.9)% |
| Height | 5961.4 (197.6) | 31.1 (1.0)% | 18.4 (2.4) | 0.9 (0.1)% |
| Body mass index (BMI) | 935.8 (198.6) | 4.9 (1.0)% | 0.4 (0.5) | 0.0 (0.0)% |
| High density lipoprotein (HDL) | 5860.8 (681.9) | 30.6 (3.6)% | 16.2 (8.3) | 0.8 (0.4)% |
| Low density lipoprotein (LDL) | 8236.4 (494.3) | 43.0 (2.6)% | 29.6 (7.8) | 1.4 (0.4)% |
| Cholesterol | 7026.0 (660.1) | 36.7 (3.4)% | 20.2 (6.8) | 0.9 (0.3)% |
| IGF1 | 3305.2 (371.8) | 17.3 (1.9)% | 4.0 (1.2) | 0.2 (0.1)% |
| Creatinine | 2052.4 (375.8) | 10.7 (2.0)% | 1.2 (1.3) | 0.1 (0.1)% |
| Red blood cell count (RBC) | 3745.8 (660.4) | 19.6 (3.4)% | 6.2 (3.6) | 0.3 (0.2)% |
| White blood cell count (WBC) | 1996.6 (120.5) | 10.4 (0.6)% | 0.6 (0.5) | 0.0 (0.0)% |
| Bone mass density in heel<br>(BMD) | 1654.2 (152.5) | 8.6 (0.8)% | 2.0 (2.3) | 0.1 (0.1)% |
| Hypertension | 257.4 (78.1) | 1.3 (0.4)% | 0.0 (0.0) | 0.0 (0.0)% |
| Cardiovascular (CVD) | 125.4 (57.7) | 0.7 (0.3)% | 0.0 (0.0) | 0.0 (0.0)% |
| **Average (s.d.)** | **4027.9 (3398.3)** | **21.0 (17.8) %** | **17.7 (35.5)** | **0.8 (1.6) %** |
| **t = 99<sup>th</sup>** |  |  |  |  |
| Hair color | 18398.6 (208.4) | 87.4 (1.0)% | 4.4 (1.5) | 2.1 (0.7)% |
| Height | 14442.6 (147.6) | 68.6 (0.7)% | 0.6 (0.9) | 0.3 (0.4)% |
| Body mass index (BMI) | 5254.4 (739.1) | 24.9 (3.5)% | 0.2 (0.4) | 0.1 (0.2)% |
| High density lipoprotein (HDL) | 14167.6 (691.4) | 67.3 (3.3)% | 0.2 (0.4) | 0.1 (0.2)% |
| Low density lipoprotein (LDL) | 15615.8 (448.1) | 74.1 (2.1)% | 0.6 (0.5) | 0.3 (0.3)% |
| Cholesterol | 14793.2 (668.3) | 70.2 (3.2)% | 0.2 (0.4) | 0.1 (0.2)% |
| IGF1 | 11049.2 (597.9) | 52.5 (2.8)% | 0.2 (0.4) | 0.1 (0.2)% |
| Creatinine | 8337.2 (702.7) | 39.6 (3.3)% | 0.0 (0.0) | 0.0 (0.0)% |
| Red blood cell count (RBC) | 11532.8<br>(1056.9) | 54.8 (5.0)% | 0.0 (0.0) | 0.0 (0.0)% |
| White blood cell count (WBC) | 8496.6 (370.7) | 40.3 (1.8)% | 0.0 (0.0) | 0.0 (0.0)% |
| Bone mass density in heel<br>(BMD) | 7816.0 (511.1) | 37.1 (2.4)% | 0.0 (0.0) | 0.0 (0.0)% |
| Hypertension | 2378.8 (390.7) | 11.3 (1.9)% | 0.0 (0.0) | 0.0 (0.0)% |
| Cardiovascular (CVD) | 1506.6 (512.3) | 7.2 (2.4)% | 0.0 (0.0) | 0.0 (0.0)% |
| **Average (s.d.)** | **10291.5 (5220.4)** | **48.9 (24.8) %** | **0.49 (1.2)** | **0.2 (0.6) %** |

For completeness, we investigated the impact of the threshold *t*, and credible level *ρ*, on PRS-based stratification uncertainty, defined as the proportion of above-threshold individuals classified as “certain above-threshold” for a given trait. As expected, the proportion of certain above-threshold classifications decreases as *ρ* increases (Figure 4a). For traits with higher average uncertainty (as defined using the scaled SD) we observe lower rates of certain classifications across all values of *ρ*. For example, at *t* =90^th^ and *ρ* =95%, the proportion of above-threshold individuals classified with certainty is 0 % for BMI (average scaled SD = 1.54) and 6.2% for hair color (average scaled SD = 0.62) (Figure 5a). Height and HDL have similar average levels of uncertainty (average scaled SD of 0.95 for height and 0.96 for HDL) and similar proportions of above-threshold individuals classified with certainty. For example, at *t* =90^th^ and *ρ* =95%, the proportions of certain classifications among above-threshold individuals are 0.9% and 0.8% for both height and HDL (Figure 5a, Table 1). Using a more stringent threshold *t* amplifies the effect of uncertainty on PRS-based stratification (Figure 5b). For example, for BMI and hair color, the proportion of certain classifications among above-threshold individuals drops for all values of *ρ* when we increase the threshold from *t*=90^th^ percentile to *t*=99^th^ percentile (Figure 5b).

We also quantified the impact of inferential variance in 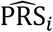 on PRS-based ranking of the test-group individuals. Using two random samples of genetic effects from one MCMC chain after burn-in, we generated two independent rankings for all individuals in the test data and quantified the correlation in the rankings (Figure 4c, Methods). We observe large variability in the rankings across the test data, with the correlation of rankings ranging from 0.25 to 0.78 across the 13 traits. We also estimated 95% credible intervals for the rank of individuals at a given percentile (e.g., 90^th^) (Table 2, Methods, Supplementary Figure 13) to find high variability in the ranking. For example, in the case of HDL an individual at 90^th^ (99^th^) percentile based on PRS point estimate can be within 41^th^ to 99^th^ percentiles (72^th^-99^th^) with 95% probability when the inferential variance in PRS estimation is taken into consideration (Table 2).

**Table 2.** Average 95% posterior ranking credible intervals for individuals at two stratification thresholds for 11 traits. We estimated the 95% posterior ranking credible intervals for individuals at the 90^th^ and 99^th^ percentiles of the testing population PRS estimates. Mean and standard deviation are calculated from the 95% posterior ranking intervals of individuals whose point estimates lie within 0.5% of the stratification threshold (213 individuals between the 89.5^th^ and 90.5^th^ percentiles for t = 90^th^ and between the 98.5^th^ and 99.5^th^ percentiles for t = 99^th^).

| Trait | t = 90<sup>th</sup> |  | t = 99<sup>th</sup> |  |
| --- | --- | --- | --- | --- |
|  | Lower bound | Upper bound | Lower bound | Upper bound |
| Hair color | 57.9 (1.8) | 97.9 (0.22) | 88.0 (2.2) | 99.8 (0.05) |
| Height | 43.4 (2.1) | 98.6 (0.18) | 74.9 (3.4) | 99.9 (0.04) |
| Body mass index (BMI) | 22.9 (2.1) | 99.0 (0.17) | 45.8 (4.0) | 99.8 (0.04) |
| High density lipoprotein (HDL) | 41.3 (2.8) | 98.7 (0.18) | 72.3 (4.1) | 99.9 (0.04) |
| Low density lipoprotein (LDL) | 49.1 (2.4) | 98.6 (0.19) | 77.7 (3.5) | 99.9 (0.04) |
| Cholesterol | 45.1 (2.8) | 98.6 (0.19) | 74.9 (3.8) | 99.9 (0.04) |
| IGF1 | 33.2 (2.4) | 98.8 (0.17) | 63.0 (4.1) | 99.9 (0.04) |
| Creatinine | 28.0 (2.4) | 98.9 (0.17) | 54.7 (4.3) | 99.9 (0.04) |
| Red blood cell count (RBC) | 34.5 (2.7) | 98.8 (0.17) | 64.4 (4.5) | 99.9 (0.04) |
| White blood cell count (WBC) | 28.2 (2.0) | 98.9 (0.17) | 56.0 (3.9) | 99.9 (0.04) |
| Bone mass density in heel (BMD) | 26.0 (2.2) | 98.9 (0.18) | 52.5 (4.1) | 99.9 (0.04) |
| Hypertension | 17.7 (1.8) | 99.0 (0.17) | 36.6 (3.4) | 99.8 (0.05) |
| Cardiovascular (CVD) | 15.5 (1.9) | 99.0 (0.18) | 32.3 (3.8) | 99.8 (0.06) |
| *Average (s.d.)* | **34.2 (12.9)** | **98.8 (.03)** | **61.0 (16.6)** | **99.9 (0)** |

### Integrating individual-PRS uncertainty into PRS-based stratification

In contrast to current PRS-based stratification practices which compare an individual’s PRS point estimate, 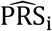, to a given threshold *t* without incorporating uncertainty, here we explore the use of the posterior probability that GV for individual *i* is above the threshold (i.e. Pr(GV_*i*_ > *t*)). We estimate Pr(GV_*i*_ > *t*) using Monte Carlo integration within the LDpred2 framework and show in simulations that the probability is well-calibrated for different causal effect size distributions despite slight miscalibration when polygenicity is high or causal variants are not present in the training SNP panel (Methods, Supplementary Figure 14 and 15). As a motivating example, two individuals with similar PRS point estimates that happen to lie on either side of a prespecified threshold (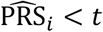 and 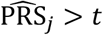) could have similar probabilities for the genetic value to exceed *t* (e.g., Pr (GV_*i*_ > *t*) = 0.4 and Pr (GV_*j*_ > *t*) = 0.6) (Figure 2).

As expected, for traits with higher PRS uncertainty, we observe a smaller proportion of testing individuals with deterministic classification (Pr(*GV*_*i*_ > *t*) = 0 or 1) (Supplementary Figure 16). We also find a tight correlation between 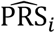 and Pr(GV_0_ > *t*) across individuals in the test data (Supplementary Figure 16). This is due to the relatively high polygenicity of the real traits in the analysis; a lower correlation is expected for traits with lower polygenicity (Supplementary Figure 17). However, Pr(GV_0_ > *t*) also contains information about individual-level false positive (FP) and false negative (FN) probabilities which, given a situation-specific cost function, can be used to calculate the expected cost of an above-threshold versus below-threshold classification (Methods). The cost functions for FP and FN should be carefully specified in the context of the clinical application. As an example, consider a scenario in which an individual’s genetic information is being used to decide whether or not to perform a bone density scan. The cost functions for FP and FN will depend on, among many other factors, the cost of a bone density scan and whether the potential benefits outweigh the risks associated with exposure to low-dose x-rays. As an example of utility of the probabilities, consider three cost functions which relate the relative costs of false positive versus false negative diagnoses: (a) equal cost for each FP and FN diagnosis (C_FP_ = C_FN_ = 1); (b) 3x higher cost for FP diagnoses (C_FP_ = 3, C_FN_ = 1); and (c) 3x higher cost for FN diagnoses (C_FP_ = 1, C_FN_ = 3). For an individual with Pr(GV_*i*_ > *t*) = 0.6, the probability of a FP versus FN diagnosis is 0.4 versus 0.6, respectively. The expected costs of FP diagnoses (Pr(FP) × C_FP_) under each scenario are (a) 0.4, (b) 1.2, and (c) 0.4; the expected costs of FN diagnoses (Pr(FN) × C_FN_) are (a) 0.6, (b) 0.6, and (c) 1.8. Therefore, the classification for this individual that minimizes the expected cost under each scenario is (a) above-threshold, (b) below-threshold, and (c) above-threshold. Assuming the same three cost functions as above, we find that the optimal decision threshold on Pr(GV_*i*_ > *t*) that maximizes the utility of the cost/gain models differs under the three functions. For C_FP_ = C_FN_ = 1, both the estimated cost curve and true cost curve achieve minimum cost at threshold = 0.5. For C_FP_ =3, C_FN_ = 1, the estimated optimum is 0.25 and the true optimum is 0.3. For C_FP_ =1, C_FN_ = 3, the estimated optimum is 0.75 and the true optimum is 0.7. More notably, assuming the probabilities are well-calibrated, we can estimate the expected cost with the individual probability of being at above-threshold, with the estimated cost curve being very close to the true cost curve despite slight inflation (Figure 7).

**Figure 6.**
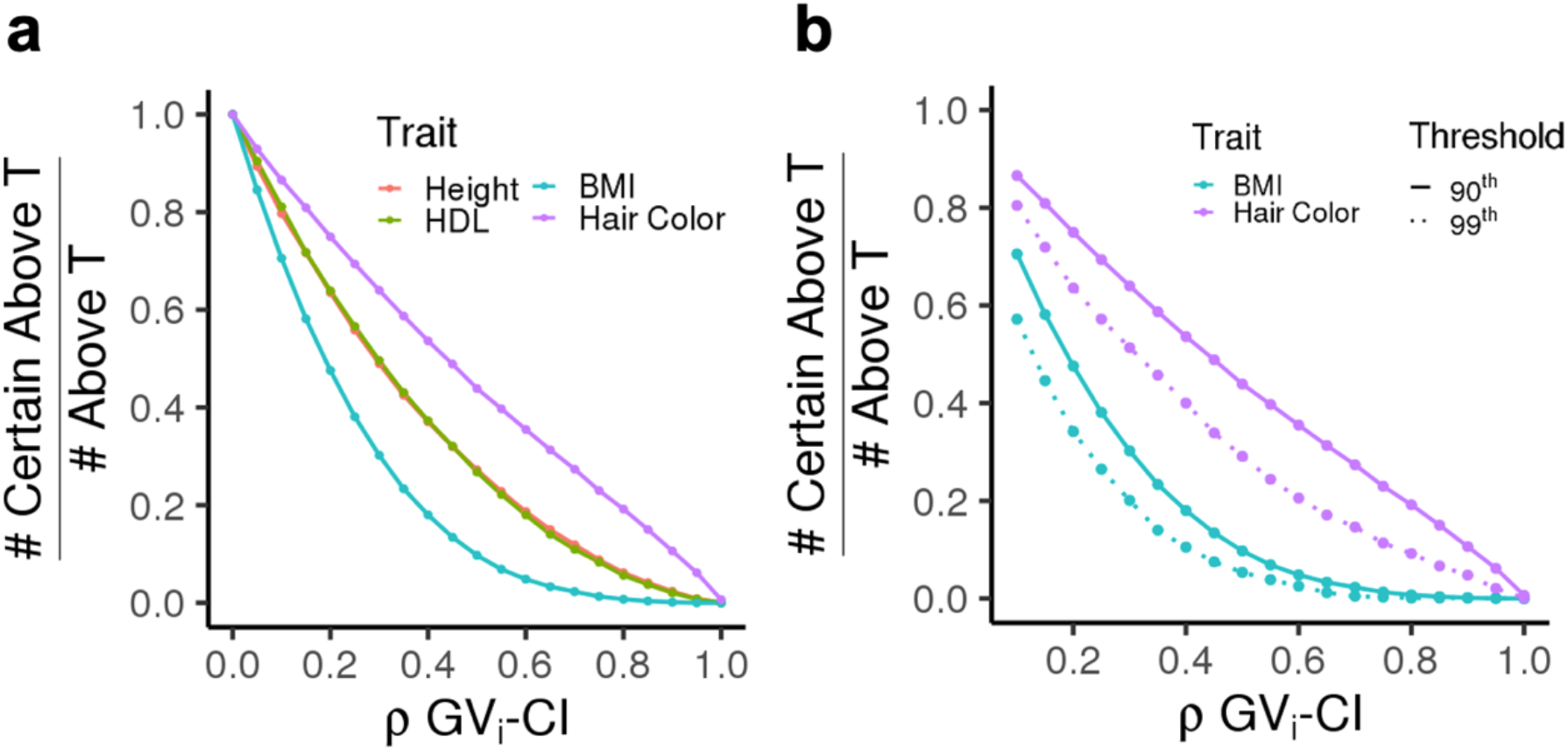
Impact of threshold *t* and credible set level *ρ* on stratification uncertainty. (a) Proportion of above-threshold classifications that are “certain” for four representative traits. The x-axis shows ***ρ*** varying from 0 to 1 in increments of 0.05. The stratification threshold *t* is fixed at 90%. (b) Proportion of above-threshold classifications that are “certain” for two representative traits and two stratification thresholds (*t* = 90%, 99%).

**Figure 7.**
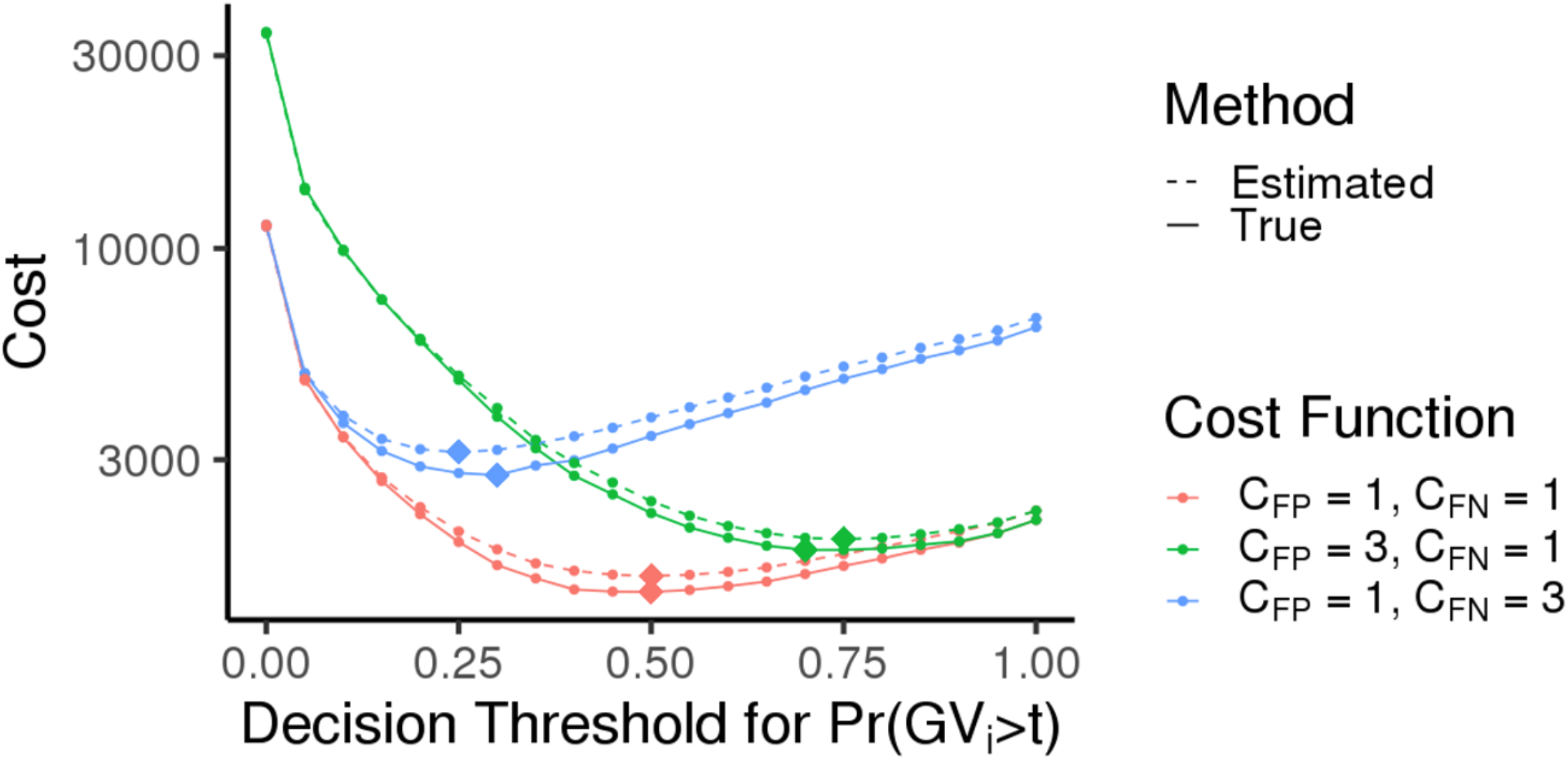
Flexible cost optimization with probabilistic individual stratification under various cost functions. Each color corresponds to one cost function: (a) equal cost for each FP and FN diagnosis (C_FP_ = C_FN_ = 1, red); (b) 3x higher cost for FP diagnoses (C_FP_ = 3, C_FN_ = 1, green); and (c) 3x higher cost for FN diagnoses (C_FP_ = 1, C_FN_ = 3, blue). The probability threshold for classification is varied along the x-axis. Solid lines represent cost calculated using true genetic risk and dotted lines represent cost estimated from the probability of an individual being above-threshold. Diamond symbols represent the optimal classification threshold for each curve (the minima). Simulation parameters are fixed to 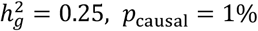.

## Discussion

In this work, we demonstrate that uncertainty in PRS estimates at the individual level can have a large impact on subsequent analyses such as PRS-based risk stratification. We note that this work focuses estimating genetic value rather than predicting phenotype; uncertainty in predictions of phenotype will be larger than the results reported here due to the additional uncertainty in unmeasured environmental factors^46^. We propose a general procedure for obtaining estimates of individual-PRS uncertainty which can be applied to a wide range of existing PRS methods. Among 13 real traits in the UK Biobank, we find that even with GWAS sample sizes on the order of hundreds of thousands of individuals, there is considerable uncertainty in individual PRS estimates (i.e. large *ρ*-level credible intervals) that can impair the reliability of PRS-based stratification. We propose a probabilistic approach to stratification that can be used in conjunction with situation-specific cost functions to help inform PRS-based decision-making, noting that such an approach is not necessarily useful for all downstream applications of PRS. Since PRS must be combined with non-genetic risk factors (e.g., age, lab values) to evaluate an individual’s absolute risk for a given disease—the quantity of interest in risk prediction—the practical utility of PRS, including measures of uncertainty in PRS, is highly dependent on disease-specific factors such as heritability, age of onset, and the costs/risks that would be incurred by initiating treatment, among many others^1,3^. Measures of uncertainty for many non-genetic risk factors are routinely propagated in risk assessment^47,48^. For example, an individual’s uncertainty-adjusted non-genetic risk factor could be one of many risk factors within a proportional hazards model^3,41,49^. We conjecture that measures of individual-PRS uncertainty will be most useful for characterizing individuals whose combined risk scores (genetics + non-genetics factors) are at or close to the decision threshold for medical intervention; we leave an investigation of uncertainty in combined risk scores for future work.

Our work is complementary to methods that aim to improve cohort-level metrics of PRS accuracy such as R^2^ or the area under the receiver operating characteristic (AUROC). We show that, for the purpose of genetic risk stratification, incorporating individual uncertainty is important as it allows us to estimate individual absolute and relative genetic risks without a validation sample, which is normally required to estimate absolute risks. As the individualized absolute risk estimates (genetic values) do not depend on a validation sample, we believe they could be robust leads to our proposed probabilistic genetic risk stratification, which can be seen as a principled approach for genetic risk stratification in clinical settings.

We conclude with several caveats and future directions. First, we quantify individual PRS uncertainty by extending LDpred2^24^, which is just one of many existing Bayesian methods that can be adapted for the same purpose (e.g., SBayesR^27^, PRS-CS^50^ and AnnoPred^51^). Extensions of other methods, including analogous procedures for P+T (PRSice-2^52^) and regularization-based approaches (lassosum^22^ and BLUP prediction^23 24^), could also be investigated. Overall, our methods produce well-calibrated credible intervals in realistic simulation parameter ranges, albeit slight mis-calibration when polygenicity is low and heritability is high. We hypothesize that it is due to several approximations employed in LDpred2 for computational efficiency. We leave investigation of the impact of approximation on calibration and further improvement for future work.

Second, while we find broad evidence that both trait-specific genetic architecture parameters (e.g., heritability, polygenicity) and individual-specific genomic features (e.g., cumulative number of effect alleles) can impact individual PRS uncertainty, both sources of uncertainty merit further exploration. For example, we perform simulations under a model in which each causal variant explains an equal portion of total SNP-heritability but, in reality, genetic architecture can vary significantly among different traits. Does individual PRS uncertainty change if both monogenic and polygenic disease risk factors^53,54^ are used for PRS estimation? We do not find a correlation between an individual’s cumulative number of effect alleles and their individual PRS uncertainty. This is primarily due to the high polygenicity of the traits being tested. Consequently, we observe tight correlation between 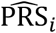 and Pr(GV_*i*_ > *t*) in most simulation scenarios except those with low polygenicity. Extending these analyses to traits with a wider range of genetic architectures will be of interest. We leave a detailed investigation of the various sources contributing to individual PRS uncertainty for ongoing work.

Third, we perform all simulations and real data analyses using genotyped SNPs (MAF > 1% on the UK Biobank Axiom Array). Since the array is designed such that the genotyped SNPs tag most of the signal from unobserved SNPs, the SNPs (predictors) used in our real data analyses likely capture most of the SNP-heritability for each trait. However, it is unclear whether individual PRS uncertainty would increase or decrease if imputed data were used instead of genotyped SNPs. Moreover, for many diseases, the largest GWAS are only available as summary statistics (estimates of marginal effects and their standard errors). It is important to assess whether there is larger uncertainty in causal effects inferred from summary statistics as that would lead to higher variability in estimated PRS. We conjecture that changes in uncertainty will also vary across traits depending on factors such as the number of SNPs (predictors) included in the PRS; the resolution of the credible sets generated by sampling causal configurations; and differences in LD tagging between predictor SNPs and causal SNPs as well as among predictor SNPs. A comparison of individual PRS uncertainty with respect to array data, imputed data, and summary statistics merits thorough investigation in future work.

Fourth, although we have shown that our approach is robust to certain types of model misspecification (e.g., effect sizes drawn from mixture of normal distributions, imperfect tagging of causal effects), we do not exclude the possibility of nonlinear interaction effects such as GxE, GxG and dominance effects^55–58^. We also assume that phenotypes are normally distributed or can be properly quantile normalized. For phenotypes with skewed distributions, the interpretation of the estimated genetic value and the associated uncertainty is unclear. For binary traits, the impact of disease prevalence and case/control sample sizes on PRS uncertainty and the interpretation of PRS uncertainty with respect to liability and odds ratio remain unclear. We leave a full investigation of these questions for future work.

Lastly, in the present study, we did not investigate individual PRS uncertainty in transethnic or admixed population settings. Causal variants, causal effect sizes, allele frequencies, and LD patterns can vary significantly across populations^59,60^. Moreover, PRS prediction accuracy (measured via cohort-level metrics) is well known to depend heavily on the ancestry of the individuals in the GWAS training data^61,62^.We therefore leave a detailed exploration of individual PRS uncertainty with respect to ancestry as future work.

## Methods

### Individual PRS uncertainty

Let *y*_i_ be a trait measured on the *i*-th individual, **x**_*i*_ an M × 1 vector of standardized genotypes and **β** an M × 1 vector of corresponding standardized effects for each genetic variant. Under a standard linear model, the phenotype model is *y* 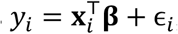, where 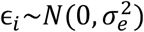. The goal of polygenic risk scores (PRS) methods is to predict genetic value for individual 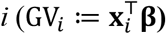 of the phenotype. In practice, the genetic effects **β** are unknown and need to be inferred from GWAS data as 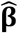. Therefore, the inferential variance in 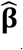 propagates to the estimated genetic value of individual *I* 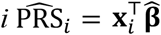. In this work we study the inferential variance in 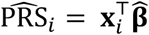 as a noisy estimate of 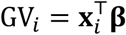.

### Estimating individual uncertainty in Bayesian models of PRS

Next, we show how Bayesian models for estimating 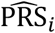 can be extended to evaluate the variance of its estimate. We focus on LDpred2, a widely used method, although similar approach can be incorporated in most Bayesian approaches. LDpred2 assumes causal effects at SNP j are drawn from a mixture distribution with spike at 0 as follows:

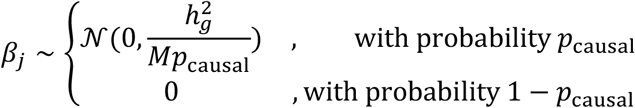

Here, *M* is the total number of SNPs in the model, 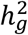 is the heritability of the trait, and *p*_causal_ is the proportion of causal variants in the model (i.e., polygenicity). Let 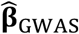 and 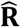 represent GWAS marginal effects and LD matrix computed from GWAS samples. By combining the prior probability 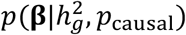 and the likelihood of observed data 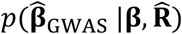, we can compute a posterior distribution as 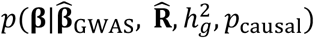. The posterior distribution is intractable and therefore LDpred2 uses Markov Chain Monte Carlo (MCMC) to obtain posterior samples from 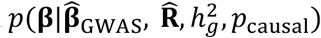 For simplicity, we use 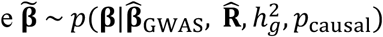 to refer to the samples from the posterior distribution, and use 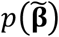 to refer to 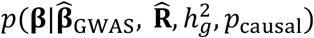 whenever context is clear. The posterior samples of the causal effects are summarized using the expectation 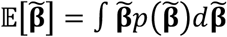, leading to 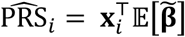.

Unlike existing methods that summarize the posterior samples of causal effects into the expectation and then estimate 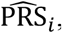, we sample from the posterior of PRS_i_ to construct a *ρ* level credible interval of genetic value (*ρ* GV_*i*_-CI) for each individual. Bernstein-von Mises theorem provides the basis that under certain conditions, such constructed Bayesian credible interval will asymptotically be of coverage probability *ρ* ^63^. This property of the Bayesian credible interval provides intuitive explanation of the uncertainty. Concretely, we obtain *B* MCMC samples from the posterior distribution of causal effects 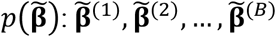. Then we compute a PRS estimate for individual *i* from each sample of 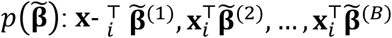 to approximate the posterior distribution of 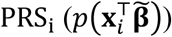. From the B samples of posterior, we obtain empirical 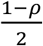 and 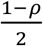 quantiles as lower and upper bound estimates of *ρ* GV_*i*_-CI (Figure 2b). As *B* goes to infinity, such Monte Carlo estimates converge to the 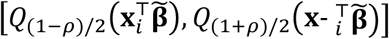, where 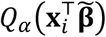 represents the *α*-quantile (here, *α* = (1 − *ρ*)/2, (1 + *ρ*)/2) for distribution of 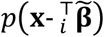 Similarly, we summarize the posterior samples using the second moment to estimate 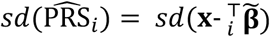. In practice, we used *B* = 500 as that leads to stable results. We investigated the autocorrelation statistics and found no evidence of autocorrelation at various lags in our experiment. (Supplementary figure 18). We recommend checking autocorrelation in practice. The MCMC samplings should be thinned when there is strong evidence of autocorrelation, which otherwise will lead to underestimation of variance.

Although in this work we focus on LDpred2, the above described procedure is generalizable to a wide range of Bayesian methods (e.g., SBayesR^27^, PRS-CS^50^ and AnnoPred^51^). Methods that are not based on Bayesian principle could potentially use Bootstrap to obtain individual uncertainty intervals^64^.

### Analytical form of individual PRS uncertainty under infinitesimal model

To facilitate understanding of PRS uncertainty, we derive an analytical estimator of PRS uncertainty under simplified assumptions: (1) all *M* SNPs are independent and causal; and (2) effect sizes are *i*.*i*.*d*. and drawn from an infinitesimal model, 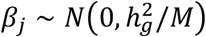 for *j* = 1,…, *M*, where 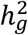 is the total heritability and *M* is the number of causal variants. Without loss of generality, we assume that genotypes are standardized to have mean zero and unit variance in the population, i.e. 𝔼 (*x*_*ij*_) = 0 and *var*(*x*_*ij*_) = 1, where *x*_*ij*_ is the genotype at SNP *j* for individual *i*. Under this assumption, following Appendix A in ref.^26^, the least squares estimate of the GWAS marginal effect 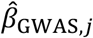 is approximately distributed as

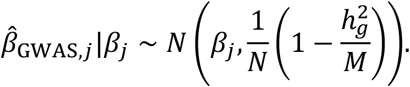

Since the per-SNP heritability in this model, 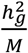, is small, the variance 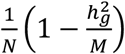 can be approximated as 1/N. The posterior distribution of 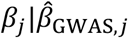 then becomes

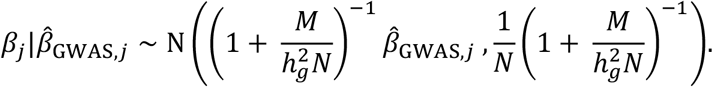

Therefore, the posterior variance of genetic value for an individual with the genotype **x**_*i*_ can be approximated as

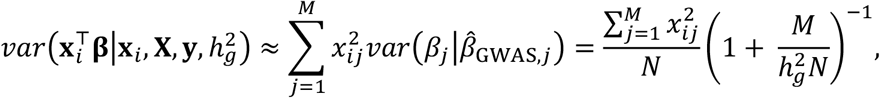

where the approximation is based on the fact that *β*_*j*_ and *β*_*k*_ are approximately independent in the posterior distribution.

Recalling that genotype is standardized so that 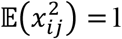, the expected posterior variance of genetic value in the population can be approximated by:

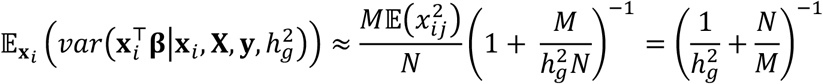

### Connection between PEV and posterior variance

Prediction error variance (PEV), a widely used concept in the animal breeding literature, is defined as 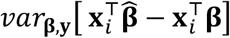, where **x**_*i*_ is the genotype of individual *i* and 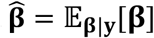 is the posterior mean of the causal effects. This variance is with respect to the randomness of both the prior **β** and phenotype **y**, holding **X** as fixed.

It follows from the law of total variance that *var*_**β**,**y**_[**β**] = 𝔼_**y**_ [*var*_**β**|**y**_[**β**]] + *var*_**y**_ [𝔼_**β**|**y**_[**β**]]. Using the fact that 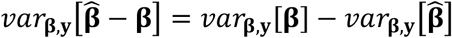 (Section 5.6.4 from ref.^31^), we have

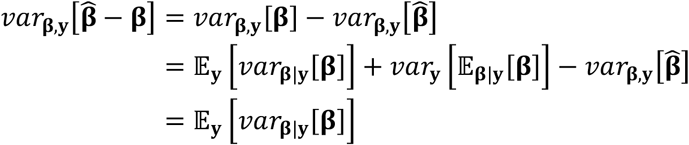

Finally, by multiplying a fixed genotype vector **x**_*i*_ to both sides, we have

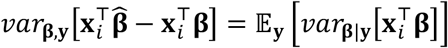

Therefore, the prediction error variance is equal to the expectation of posterior variance under repeated sampling of **y**. Given large sample sizes, we expect that for each realization of 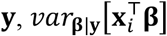 will not deviate much from 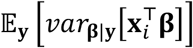. Therefore, PEV and posterior variance will be approximately equal. We also note that under infinitesimal model setting, the posterior variance of genetic value has the same matrix form as the inversion of coefficient matrix of mixed model equation for BLUP^30,33^.

### Simulations

We design simulation experiments in various settings and different sample sizes to understand the properties of uncertainty in PRS estimates. We used simulation starting from genotypes in UK Biobank ^65^. We excluded SNPs with MAF < 0.01 and genotype missingness > 0.01, and those SNPs that fail the Hardy-Weinberg test at significance threshold 10^−7^, which leaves us 459,792 SNPs. We preserve “white British individual”, with self-reported British white ancestry and filter pairs of individuals with kinship coefficient < 1/2^(9/2)^) ^65^. We further filtered individuals who are outliers for genotype heterozygosity and/or missingness, and obtained 291,273 individuals for all analyses.

Given the genotype matrix **X**, heritability 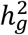, proportion of causal variants *p*_causal_, standardized effects and phenotypes are generated as follows

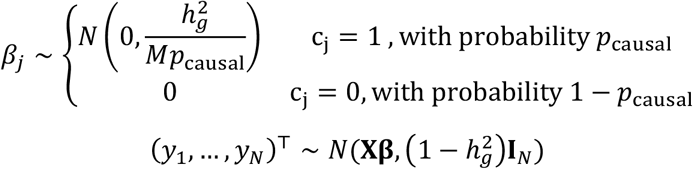

Finally, given the phenotypes **y** = (*y*_1_, …, *y*_*N*_)^T^ and genotypes **X**, we simulate the GWAS marginal association statistics with 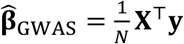. We simulate the data using a wide range of parameters, 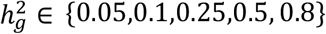, *p*_causal_ ∈ {0.001,0.01,0.1,1}, a total of 20 simulation settings, with each repeated 10 times. The total population of individuals is randomly assigned to 250,000 individuals as the training population, 20,000 individuals as the validating population, and the rest of 21,273 individuals as the testing population, as the usual practice for the PRS model building process. When investigating how sample sizes in the training cohort change PRS uncertainty, we vary the sample sizes in the training population in 20,000, 50,000, 100,000, 150,000, and 250,000, while holding the validation population and testing population as intact, to enable a fair comparison between sample sizes.

### Real data analysis

We performed real data analysis with 13 real traits from UK Biobank, including hair color, height, body mass index (BMI), bone mass density in the heel (BMD), high density lipoprotein (HDL), low density lipoprotein (LDL), cholesterol, igf1, creatinine, red blood cell count (RBC) and white blood cell count (WBC), hypertension and cardiovascular disease. The genotype was processed in the same way as the simulation study, where we have 459,792 SNPs and 291,273 individuals. We randomly partitioned the total of 291,273 individuals into 250,000 training, 20,000 validation and 21,273 testing groups. The random partition was repeated five times to average of the randomness of results due to sample partition. For each round of random partition of the individuals, we calculated marginal association statistics between genotype and quantile-normalized phenotype in training group with PLINK, using age, sex, and the first 20 genetic principal components as the covariates. Then we applied LDpred2 to obtain the individual posterior distribution of the genetic value, as described above. We regressed out covariates from the phenotypes to obtain adjusted phenotypes, where the regressing coefficients are first estimated from the training population, and applied to phenotype from training, validation and testing population respectively. We evaluate accuracy of PRS estimates in validation and testing groups by Pearson correlation between PRS estimates and adjusted phenotypes.

### PRS analysis using LDpred2

We run LDpred2 for both simulation and real data analysis with the following settings. We calculate the in-sample LD with functions provided by the LDpred2 package, using the window size parameter of 3cM. We estimate the heritability 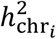, *i* = 1, …, 22 for each chromosome with built-in constrained LD score regression^66^ function. We run LDpred2-grid per chromosome with a grid of 17 polygenicity parameters *p*_causal_ from 10^−4^ to 1 equally spaced in log space, three heritability Parameters 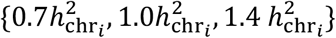, and with the sparsity option both enabled and disabled, as recommended by LDpred2. We choose the model with the highest R^2^ between the predicted posterior mean and the (adjusted) phenotype on validation set as best model to apply to testing data. We extract 500 posterior samples of causal effects 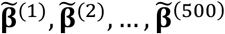 after 100 burn-in iterations from MCMC sampler of the model to approximate posterior distribution of causal effects. For each individual with genotype **x**_*i*_, we calculate 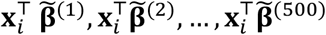 to approximate GV posterior distribution for individual *i*. We then calculate summary statistics of GV posterior distribution, including the posterior mean 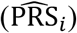, *ρ* level credible interval (*ρ* GV_*i*_-CI) and probability of above threshold t (Pr(GV_*i*_ > t)).

### Calculating and evaluating the coverage

We evaluate the coverage properties of *ρ* GV_i_-CI in simulation: we check whether 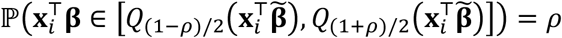. To evaluate this property, for each simulated dataset, we calculate the frequency of the true genetic risk lies in the predicted interval, i.e., the frequency of 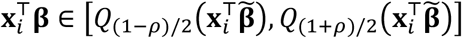 for every individual in the testing population, for *ρ* ∈ {0.1, 0.2, …, 1.0}. This property provides us an intuitive understanding of the predicted interval: for an individual with a predicted interval 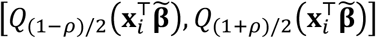, its true genetic risk is expected to be in this interval with a probability *ρ*.

### Definition of scaled standard deviation in individual PRS estimates

To compare the relative order of standard deviation across different genetic architecture, especially across genetic architecture with different heritability, we define the quantity, scaled standard deviation in individual PRS estimates (scaled 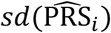) to enable fair comparison. The quantity is defined for every individual *i*, as 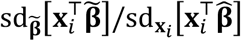, where the numerator term 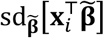 refers to standard deviation due to the posterior sampling of 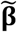 of *i* -th individual. Recalling that 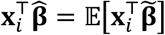, the denominator term 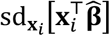 refers to the variation of the point estimate across individuals in the population.

### Posterior individual ranking interval

The relative rank of individual 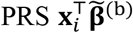 in the population 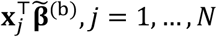 varies across different MCMC samplings of posterior causal effects. To evaluate the uncertainty of ranking for individual *i*, we compute 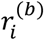 as the quantile of 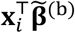 in the population 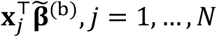 for each of the *b* = 1, …, *B* posterior samples to approximate posterior distribution of the relative rank. We can obtain ρ-level credible intervals of ranking as [*Q*_(1−*ρ*)/2_(*r*_*i*_), *Q*_(1+*ρ*)/2_(*r*_*i*_)] for each individual *i*. To assess the uncertainty of ranking for individuals at 90 (99) percentile threshold based on PRS estimates, we select individuals within 1 percentile of thresholds (89.5-90.5%, 98.5-99.5%) and compute mean and standard deviation for lower and upper bound of ρ=95% posterior ranking interval, across the selected individuals.

### PRS rank correlation between different MCMC samplings

With the *B* posterior causal effects samples 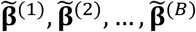 after burn-in, and *N* individuals in the testing population **x**_1_, **x**_2_, …, **x**_*N*_, we compute PRS for each individual, 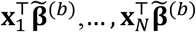 and its relative rank in the population ^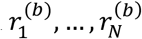^ for each posterior sample 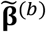. Then for each pair of different *b*_1_-th,*b*_2_-th posterior samples, 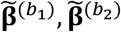, we calculate the spearman correlation between 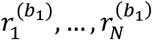 and 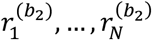, representing the variability of the ranks across MCMC samplings. We compute the rank correlation for 1000 pairs of different MCMC samplings, and get the distribution of the rank correlation.

### Probabilistic risk stratification

We define the notion of probabilistic framework for risk stratification based on posterior distribution of GV_*i*_. Given a pre-specified threshold *t*, for every individual, we can calculate the posterior probability of the genetic risk larger than the given threshold *t*, Pr(GV_i_ > *t*), with Monte Carlo integration as

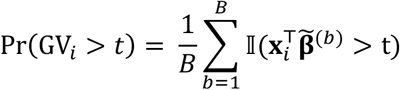

We use the previous simulation settings to show that this probability is well calibrated. For each simulation, we divide the individuals based on their posterior probability of being at above-threshold into 10 bins with {0, 0.1, …, 1.0} as breaks. For each bin, we calculate the proportion of individuals with true genetic risk higher than the threshold as the empirical probability and the average posterior probability as theoretical probability. The empirical probability is expected to be the same as theoretical probability.

### Utility analysis

The individualized posterior distribution of genetic value provides extra information for patient stratification. We consider a scenario that there is a cost associated for decision that (1) classify an individual with low genetic risk into a high genetic risk category, *C*_FP_, where FP represents false positive. (2) classify an individual with high genetic risk into a low genetic risk category, *C*_FN_, where FN represents false negative. For an individual with posterior probability Pr(GV_*i*_ > *t*), we want to decide an action, whether to classify this individual to be at high genetic risk, and perform further screening. If we classify this individual as above-threshold, we will have probability 1 − Pr(GV_*i*_ > *t*), that this individual is in fact below-threshold, inducing an expected cost *C*_*FN*_(1 − Pr(GV_*i*_ > *t*)). Conversely, if we classify this individual as below-threshold, we will have probability Pr(GV_*i*_ > *t*) that this individual will be in the high genetic risk, inducing an expected cost *C*_*FN*_ Pr(GV_*i*_ > *t*). To minimize the expected cost, we would decide according to which action leads to the least cost. The critical value in this scenario is 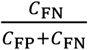: if 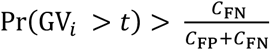, we would choose to classify this individual as above-threshold, otherwise below-threshold.

### Software implementation

Our method is implemented in the LDpred2 package (see URLs). In the function ‘snp_ldpred2_grid’, setting the option ‘return_sampling_betas = TRUE’ will output B posterior samples of the causal genetic effects. Posterior samples of an individual’s GV are obtained by multiplying the individual’s genotype by the M x B weight matrix. One can subsequently obtain the posterior mean, posterior variance, and other quantities of interest from the posterior of the GV. We note that the time required to estimate the causal effects remains the same; the only additional computational costs come from storing the M x B weight matrix and from multiplying the genotype vector by an M x B matrix rather than an M x 1 vector. The memory required to store 500 samples of causal effects for 459,792 SNPs is approximately 2 GB. Given the B posterior samples of causal effects, the runtime for computing the posterior distribution of genetic value for 10,000 testing individuals is less than five minutes.

## Data availability

The individual-level genotype and phenotype data are available by application from the UKBB http://www.ukbiobank.ac.uk/.

## URLs

LDpred2 software implementing individual PRS credible intervals: https://privefl.github.io/bigsnpr/articles/prs_uncertainty.html

Scripts for simulations and real data analyses: https://github.com/bogdanlab/prs-uncertainty

## Acknowledgments

This research was conducted using the UK Biobank Resource under application 33297. We thank the participants of UK Biobank for making this work possible. This work was funded in part by NIH awards R01HG009120, R01MH115676. R01HG006399, and U01CA194393. The content is solely the responsibility of the authors and does not necessarily represent the official views of the NIH.

