## Supplementary Tables and Figures for "Large uncertainty in individual PRS estimation impacts PRS-based risk stratification"

| Trait | $h_g^2$ | $p_{causal}$ | accuracy ($R_{phe}^2$) | average $sd(\widehat{PRS}_i)$ | average scaled $sd(\widehat{PRS}_i)$ |
|---|---|---|---|---|---|
| Hair color | 0.29 (0.02) | 0.03 (0.02) | 0.19 (0.01) | 0.27 (0.01) | 0.62 (0.02) |
| Height | 0.52 (0.04) | 0.09 (0.01) | 0.32 (0.00) | 0.40 (0.01) | 0.95 (0.03) |
| Body mass index (BMI) | 0.24 (0.01) | 0.22 (0.03) | 0.10 (0.01) | 0.46 (0.02) | 1.54 (0.06) |
| High density lipoprotein (HDL) | 0.30 (0.01) | 0.09 (0.02) | 0.18 (0.00) | 0.36 (0.02) | 0.96 (0.05) |
| Low density lipoprotein (LDL) | 0.18 (0.01) | 0.03 (0.01) | 0.10 (0.01) | 0.23 (0.01) | 0.72 (0.03) |
| Cholesterol | 0.17 (0.02) | 0.03 (0.01) | 0.09 (0.01) | 0.25 (0.01) | 0.84 (0.05) |
| IGF1 | 0.25 (0.01) | 0.06 (0.01) | 0.13 (0.01) | 0.40 (0.01) | 1.18 (0.04) |
| Creatinine | 0.21 (0.01) | 0.07 (0.01) | 0.10 (0.00) | 0.33 (0.01) | 1.33 (0.05) |
| Red blood cell count (RBC) | 0.24 (0.02) | 0.05 (0.01) | 0.13 (0.00) | 0.34 (0.02) | 1.13 (0.07) |
| White blood cell count (WBC) | 0.21 (0.01) | 0.07 (0.02) | 0.10 (0.00) | 0.40 (0.01) | 1.33 (0.03) |
| Bone mass density in heel (BMD) | 0.36 (0.02) | 0.04 (0.01) | 0.12 (0.01) | 0.45 (0.02) | 1.38 (0.04) |
| Hypertension | 0.14 (0.00) | 0.15 (0.03) | 0.04 (0.00) | 0.35 (0.01) | 1.83 (0.06) |
| Cardiovascular (CVD) | 0.14 (0.01) | 0.13 (0.05) | 0.03 (0.00) | 0.32 (0.02) | 1.97 (0.10) |

**Supplementary Table 1. Accuracy and individual PRS uncertainty in testing group.** $h_g^2$ is estimated as the sum of optimal heritability parameter in LDpred2 of 22 chromosomes. $p_{causal}$ is estimated as the sum of optimal sparsity parameter weighted by the number of SNPs of 22 chromosomes. Accuracy is measured as the Pearson correlation between PRS estimates and adjusted phenotypes. Both mean and standard deviation of $h_g^2$, $p_{causal}$ and $R_{phe}^2$ of five random data partition replicates are reported. Individual PRS uncertainty are reported as mean and standard deviation of $sd(\widehat{PRS}_i)$ and scaled $sd(\widehat{PRS}_i)$ across 21,273 individuals and five random data partition replicates (106,365 data points).

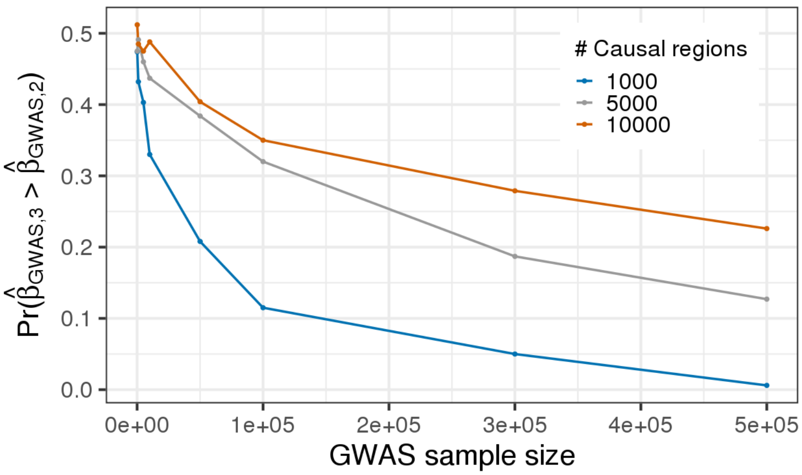


**Supplementary Figure 1. GWAS sample size and causal effect size impact the relative ordering of marginal GWAS effects at tag versus true causal SNPs.** We simulated a GWAS of $N$ individuals ($\mathbf{X_{N\times 3}}$) for 3 SNPs with LD structure $\mathbf{R}$ (SNP2 and SNP3 are in LD of 0.9 whereas SNP1 is uncorrelated to other SNPs) where SNP1 and SNP2 are causal with the same effect size $\boldsymbol{\beta_c} = (\beta, \beta, 0)$ such that the variance explained by this region is $var(\mathbf{X}\boldsymbol{\beta_c}) = 0.5/m_{causal}$ corresponding to a trait with total heritability of 0.5 equally distributed across $m_{causal}$ regions in the genome. For each parameter setting we quantified the proportion of times the marginal GWAS effect at SNP3 (tag SNP) is larger than the observed marginal effect at SNP2 (true causal) across 1,000 randomly drawn GWASs. To explore the impact of different causal effect sizes, we varied $m_{causal}$ from 1,000 to 10,000 causal regions in the genome.

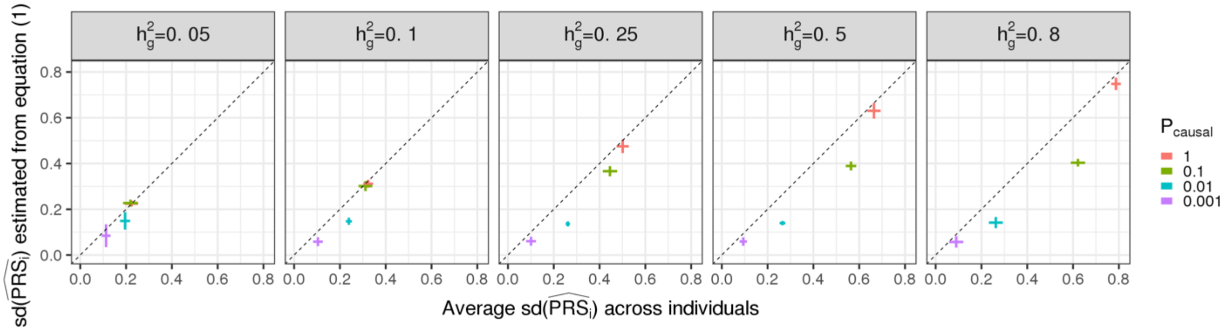


**Supplementary Figure 2. Analytical estimator of $\mathbf{sd}(\widehat{\mathbf{PRS}}_\mathbf{i})$ provides an approximately unbiased estimates of average sd(PRSi) of testing individuals.** The x-axis is the average $\mathrm{sd}(\widehat{\mathrm{PRS}}_\mathrm{i})$ in testing individuals within each simulation replicate. The y-axis is the expected $\mathrm{sd}(\widehat{\mathrm{PRS}}_\mathrm{i})$ computed with Equation (1), replacing M and $h_g^2$ with estimates of the number of causal variants and SNP-heritability, respectively, from LDpred2. Each dot is an average of 10 simulation replicates for each $\mathrm{p_{causal}} \in \{0.001, 0.01, 0.1, 1\}$. The horizontal whiskers represent $\pm$1.96 standard deviations of average $\mathrm{sd}(\widehat{\mathrm{PRS}}_\mathrm{i})$. The vertical whiskers represent $\pm$1.96 standard deviations of expected $\mathrm{sd}(\widehat{\mathrm{PRS}}_\mathrm{i})$. Note that when $p_{\mathrm{causal}} = 1$, the independent LD assumption is violated but the analytical form still provides approximately unbiased estimates. When $p_{\mathrm{causal}} \neq 1$, the infinitesimal assumption is violated, leading to downward bias in the analytical estimator. In these scenarios, since we simply replace $M$ with $M \times p_{\mathrm{causal}}$, the uncertainty identifying the causal variants is ignored by Equation (1).

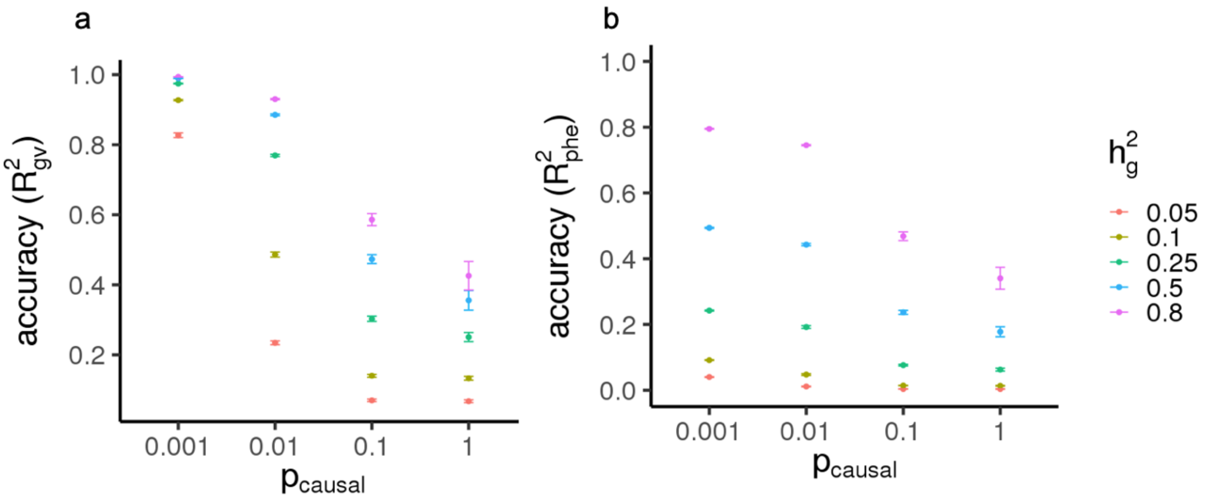


**Supplementary Figure 3. Accuracy of LDpred2 PRS estimates for testing individuals in simulation.** The accuracy is measured by two metrics: a) $R^2_{\mathrm{GV}}$, the squared Pearson correlation between PRS estimates and true genetic value. b) $R^2_{phe}$, the squared Pearson correlation between PRS estimates and phenotype. We simulate 20 parameter settings with $h^2_g \in \{0.05, 0.1, 0.25, 0.5, 0.8\}$ and $p_{causal} \in \{0.001, 0.01, 0.1, 1\}$. Different colors represent different $h^2_g$. Each point represents average accuracy of 10 repeats for each simulation settings. The error bars are $\pm$ 1.96 standard error of the mean.

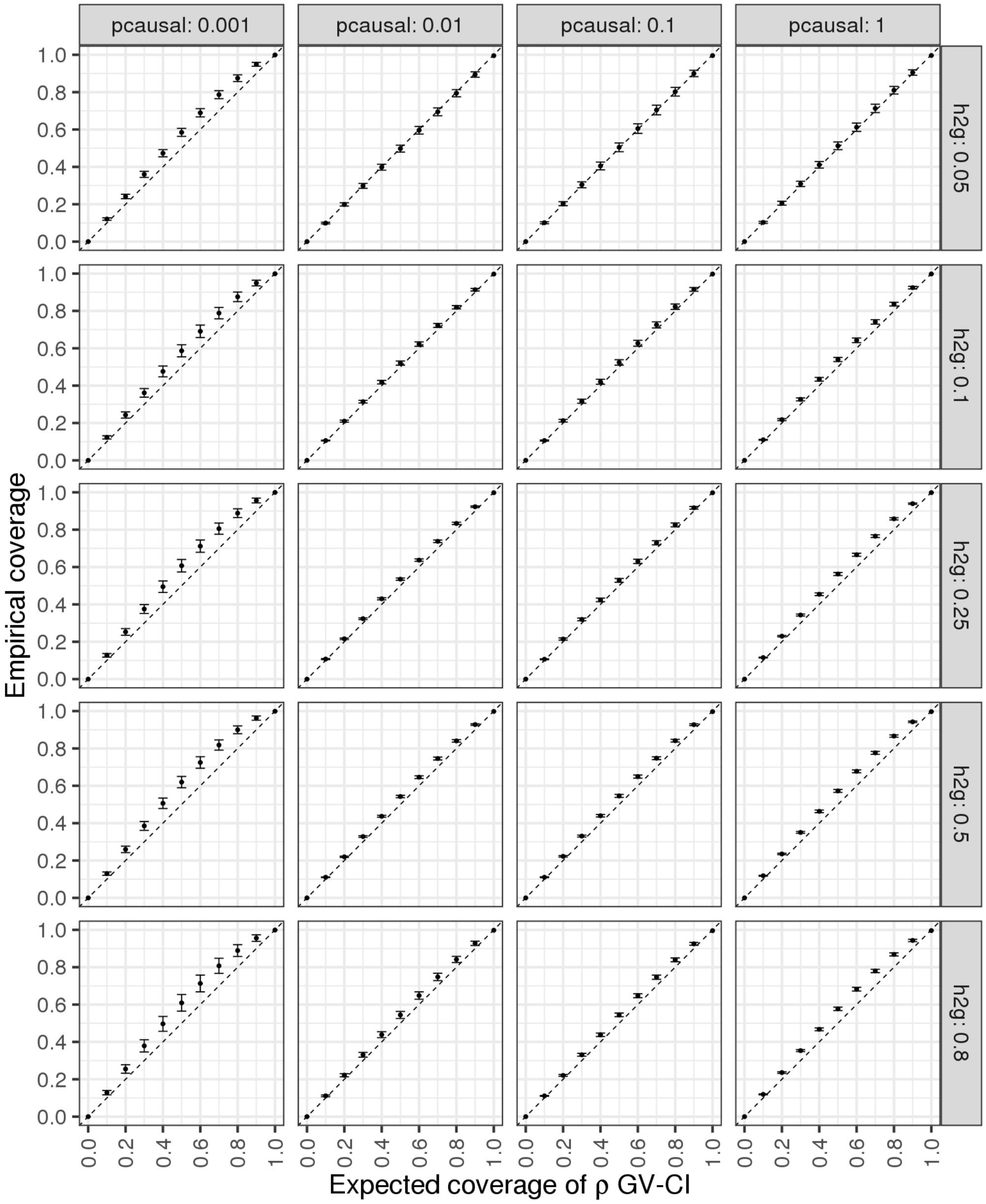


**Supplementary Figure 4. Calibration of $\rho$-level genetic value credible interval with respect to proportion of causal effects and SNP-heritability in testing individuals**. Each row of panels corresponds to one heritability parameter $h_g^2 \in \{0.05, 0.1, 0.25, 0.5, 0.8\}$ and each column of panels corresponds to one polygenicity parameter $p_{causal} \in \{0.001, 0.01, 0.1\ 1\}$. The x-axis is the expected coverage of $\rho$-GV CI ($\rho$). The y-axis is the empirical coverage calculated as the proportion of $\rho$-GV CIs that contain the true genetic value for one simulation repeat. The dot is the average empirical coverage calculated from 10 simulation repeats. The error bars are $\pm$1.96 standard error of mean.

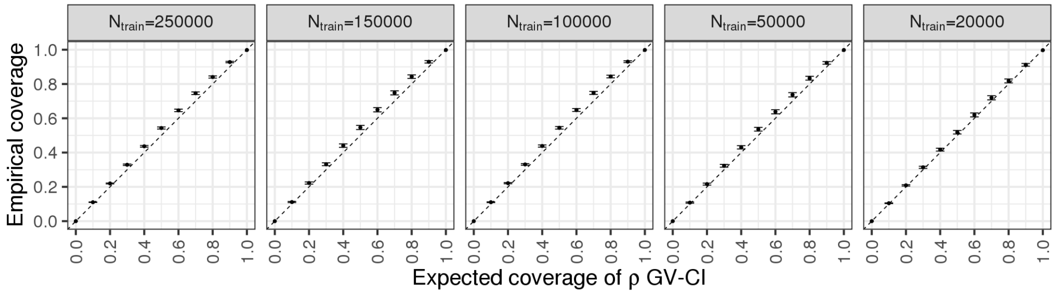


**Supplementary Figure 5, Calibration of ρ-level genetic value credible interval at various sample sizes.** Each panel represents the results with training cohort sample sizes of 250K, 150K, 100K, 50K and 20K. The x-axis is the expected coverage of ρ-GV CI (ρ). The y-axis is the empirical coverage, calculated as the proportion of ρ-GV CIs that contain the true genetic value for one simulation replicate. In all panels, h2g = 0.5, pcausal = 0.01, and the simulation contains the genome-wide UKBB array SNPs.

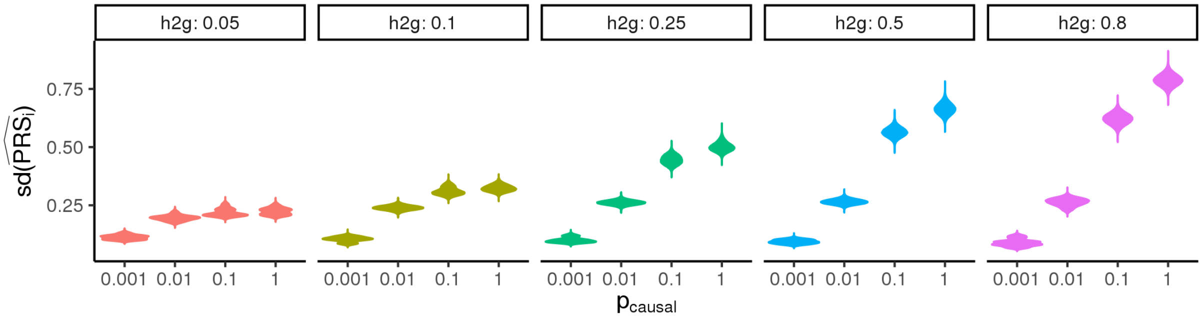


**Supplementary Figure 6. Distribution of individual PRS absolute standard deviation with respect to polygenicity under different heritability.** Each panel represents simulation with one $h_g^2$ from $\{0.05, 0.1, 0.25, 0.5, 0.8\}$. The x-axis is four polygenicity parameters ($p_{causal} \in \{0.0001, 0.01, 0.1, 1\}$). The y-axis is standard deviation in PRS estimation of an individual. Each violin plot represents 21,273 testing individuals across 10 simulations (212,730 values).

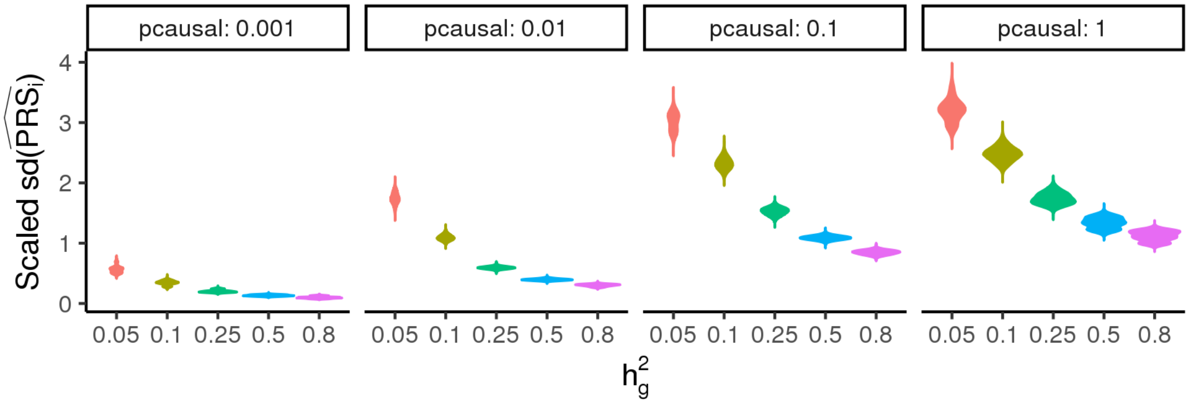


**Supplementary Figure 7. Distribution of individual PRS absolute standard deviation with respect to heritability under different polygenicity.** Each panel represents simulation with one polygenicity from $\{0.001, 0.01, 0.1, 1\}$. The x-axis is five heritability parameters ($h_g^2 \in \{0.05, 0.1, 0.25, 0.5, 0.8\}$). The y-axis is scaled standard deviation in PRS estimation of an individual. Each violin plot represents 21,273 testing individuals across 10 simulations (212,730 values).

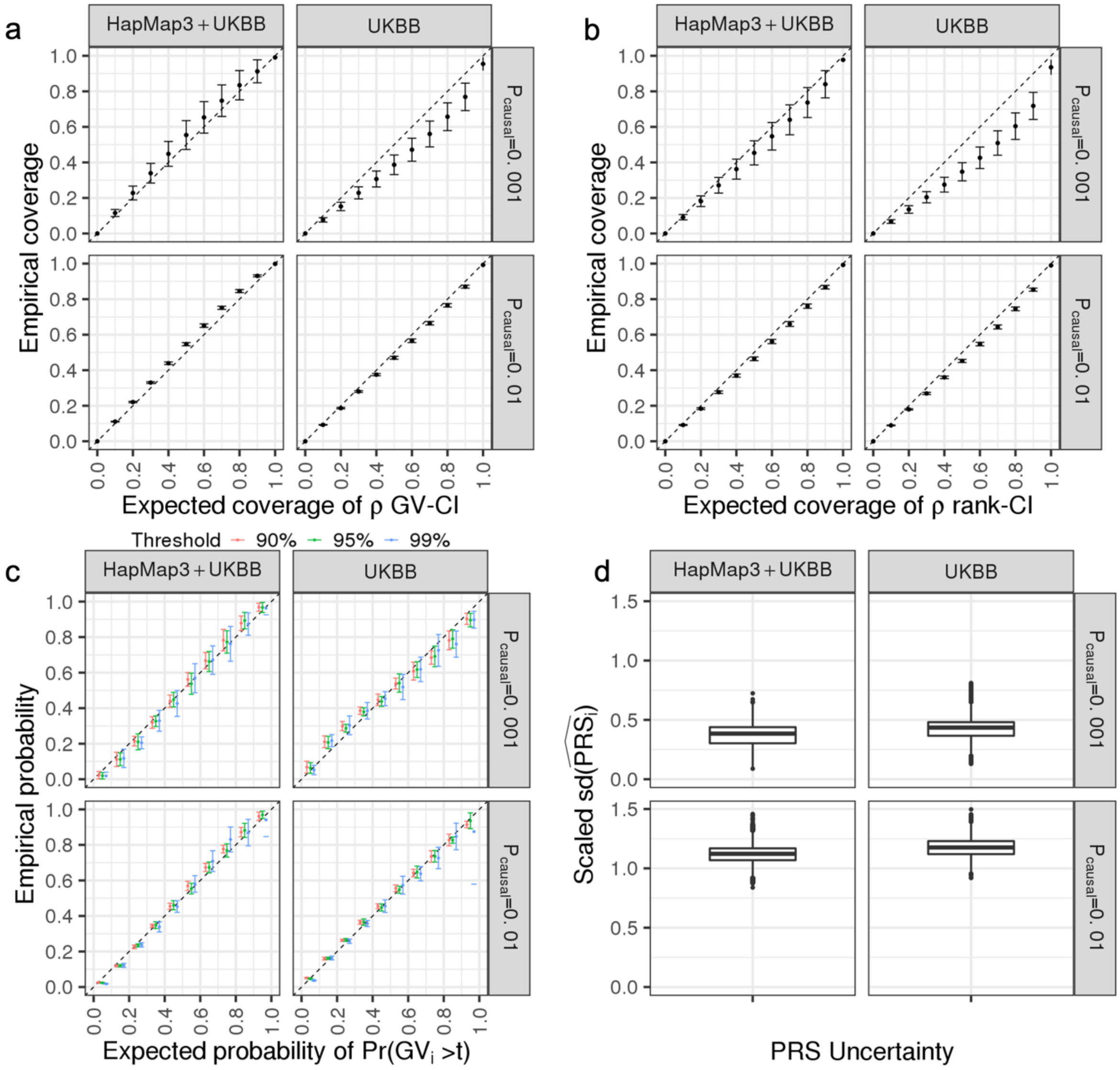


**Supplementary Figure 8. Posterior distribution of genetic value is mis-calibrated when causal variants are partially absent in the SNP panel used for PRS training.** For all panels, we simulated 124,080 SNPs (a union of 36,987 UK Biobank (UKBB) array SNPs and 93,767 HapMap3 SNPs) on chromosome 2. We trained the PRS model on either the HapMap3 + UKBB SNPs (all causal variants are observed in the training data) or UKBB SNP panel (~70% of causal variants are excluded). (a) Calibration of $\rho$-level genetic value credible interval. The x-axis is the expected coverage of $\rho$-GV CI (i.e. $\rho$). The y-axis is the empirical coverage calculated as the proportion of GV CIs that contain the true genetic value in one simulation replicate. (b) Calibration of $\rho$-level rank credible interval. The x-axis is the expected coverage of the rank CI ($\rho$). The y-axis is the empirical coverage calculated as the proportion of $\rho$-rank CIs that contain the true rank of individual among testing individuals in one simulation replicate. (c) Calibration of probability of GV above threshold t. The x-axis is the expected probability set as middle of each bin. The y-axis is the empirical probability calculated as the proportion of individuals having GV within the lower and upper bound of the bin of one simulation replicate. Different colors represent different prespecified thresholds. (d) Distribution of individual PRS scaled standard deviation. For (a-c), the dot is the average empirical coverage calculated from 10 simulation replicates. The error bars are $\pm$1.96 standard error of mean. For (d), the boxplot center line is the median; the lower and upper hinges correspond to the first and third quartiles, and boxplot whiskers extend to the minimum and maximum estimates located within 1.5 × interquartile range (IQR) from the first and third quartiles, respectively.

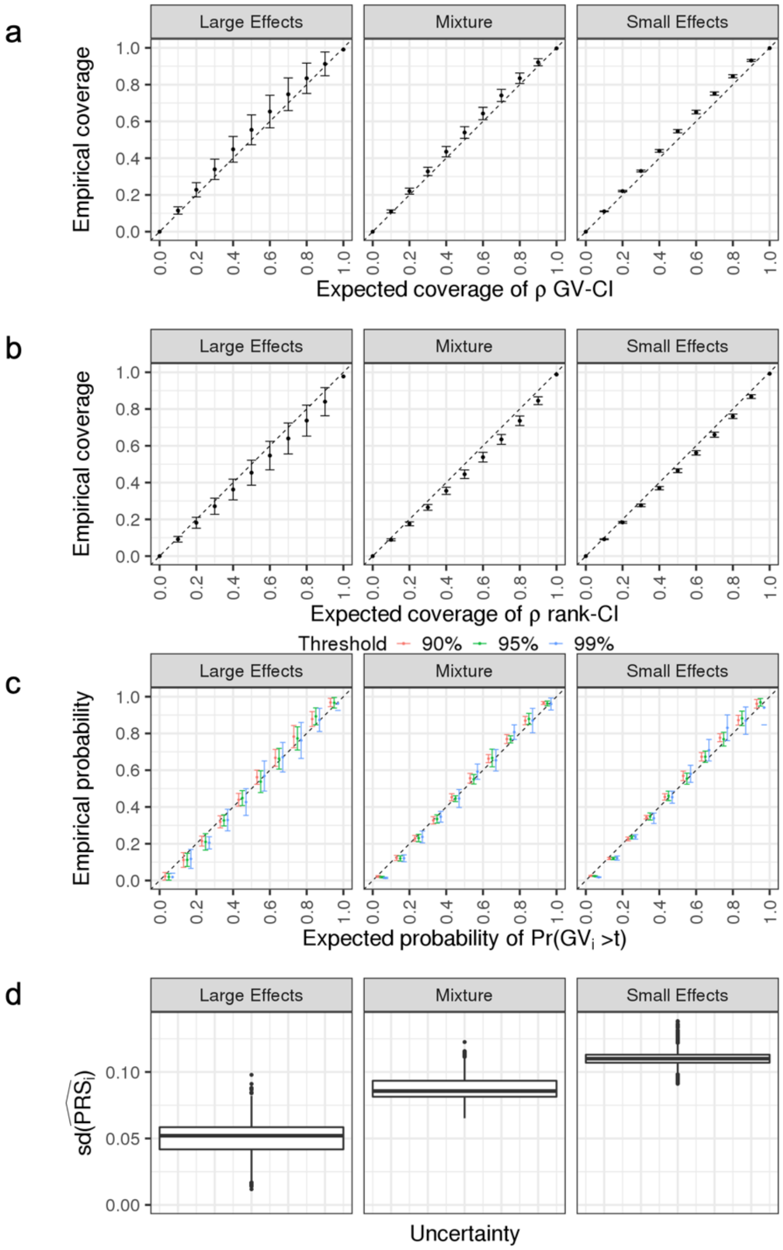


**Supplementary Figure 9. Posterior distribution of genetic value is well-calibrated for mixture of normal effect size distribution.** Each column summarizes results for each of the three genetic architectures. Small effects are simulated under pcausal = 0.01, h2g = 0.02; large effects are simulated under pcausal = 0.001, h2g = 0.02; Mixture refers to a half and half mixture of the two simulations (small effects: pcausal = 0.0005, h2g = 0.01; large effects: pcausal = 0.005, h2g = 0.01). (a) Calibration of $\rho$-level genetic value credible intervals. (b) Calibration of $\rho$-level rank credible intervals. (c) Calibration of probability of GV above threshold t. (d) Distribution of individual PRS standard deviations. See Supplementary Figure 8 for a detailed figure description.

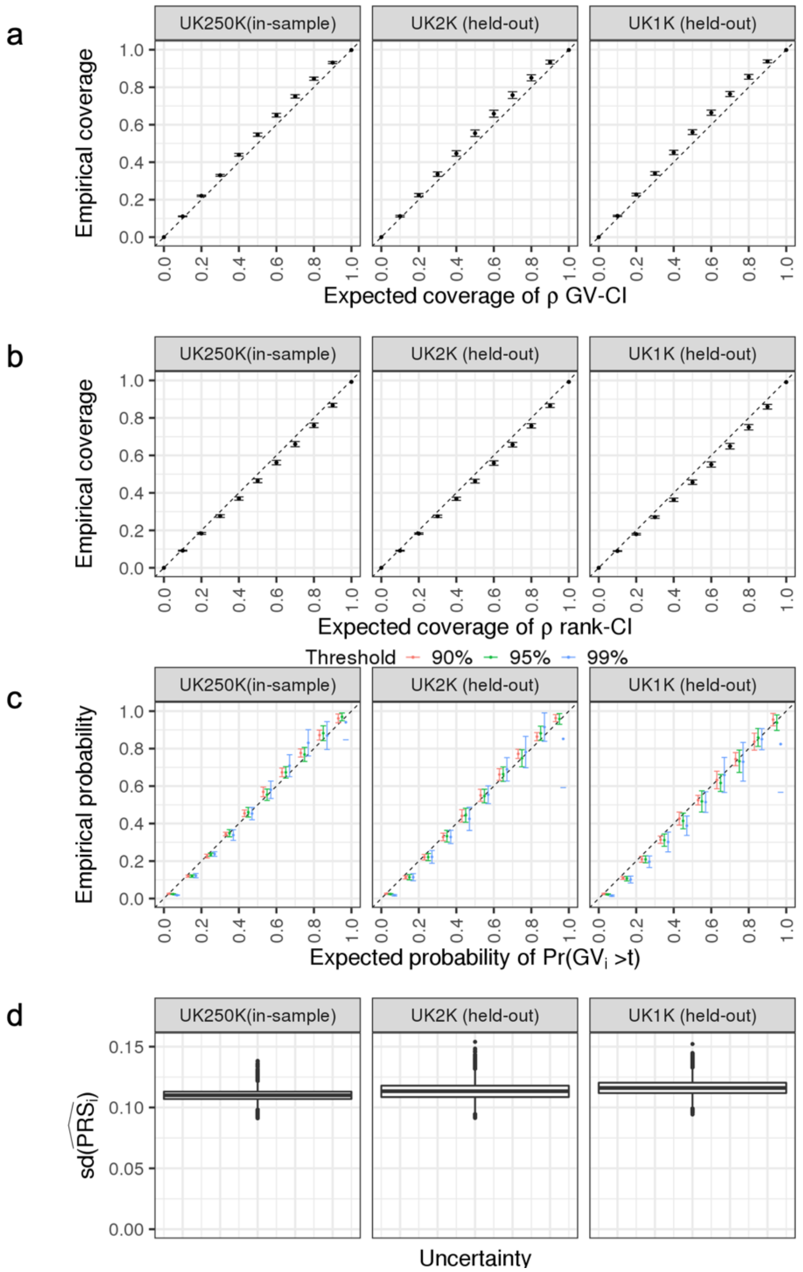


**Supplementary Figure 10. Posterior distribution of genetic value is well-calibrated with external LD.** Each column summarizes the calibration and uncertainty of PRS trained with LD computed from four different cohorts: I. 250K UKB training individuals; II. 2K held-out UKBB individuals; III. 1K held-out UKBB individuals. (a) Calibration of $\rho$-level genetic value credible intervals. (b) Calibration of $\rho$-level rank credible intervals. (c) Calibration of probability of GV above threshold t. (d) Distribution of individual PRS scaled standard deviation. See Supplementary Figure 8 for a detailed figure description ($h^2_g$ = 0.02, $p_{causal}$= 0.01).

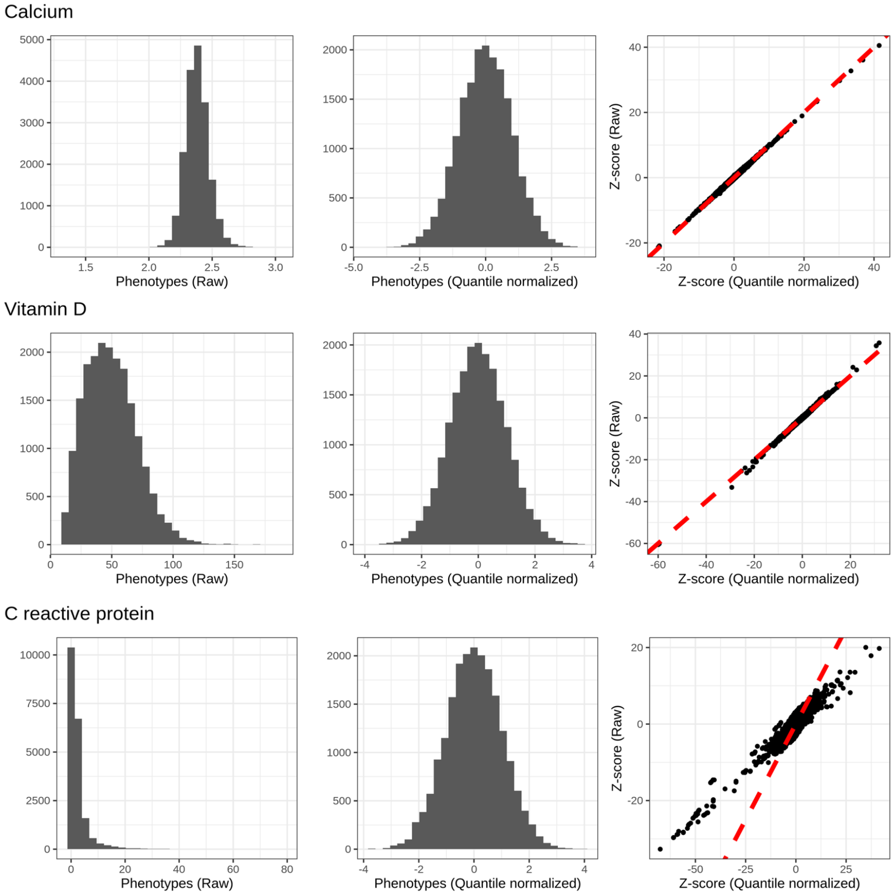


**Supplementary Figure 11. Histograms of raw phenotypes, quantile-normalized phenotypes and the corresponding consistency of the GWAS z-score.** We show the histograms of (left) phenotype distribution before quantile normalization, (middle) phenotype distribution after quantile normalization, and (right) the consistency between the marginal association z-scores calculated from phenotypes before and after quantile normalization.

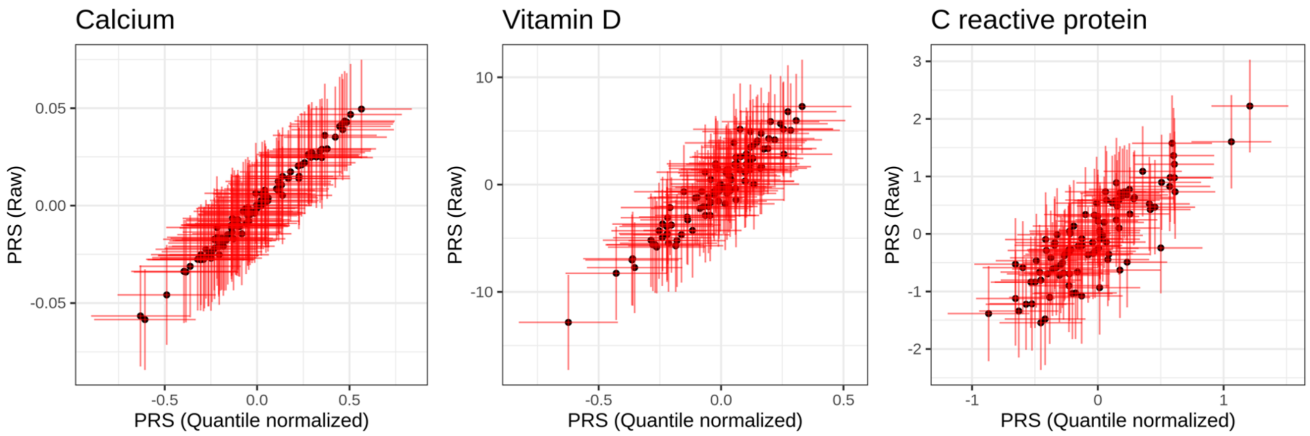


**Supplementary Figure 12. Consistency of PRS point estimates and scaled PRS SD with respect to phenotype quantile normalization.** The black dots are PRS point estimates for the quantile normalized phenotype and raw phenotype. The corresponding sd(PRSi) are marked by the red error bars.

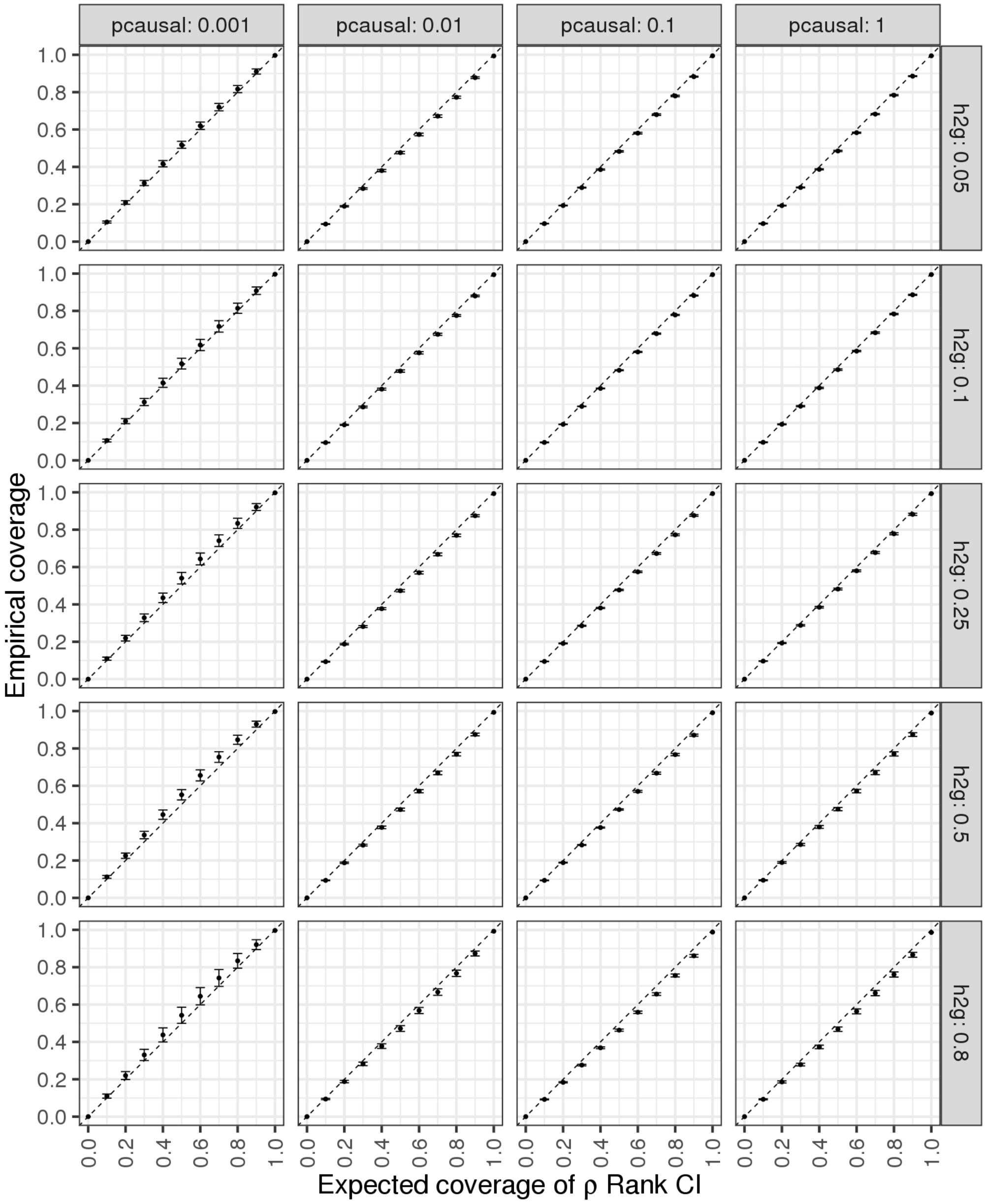


**Supplementary Figure 13. Calibration of $\rho$-level rank credible interval with respect to proportion of causal effects and SNP-heritability in testing individuals**. The x-axis is the expected coverage of $\rho$-Rank CI. The y-axis is the empirical coverage calculated as the proportion of $\rho$-Rank CIs that contain the true rank of individual among testing individuals for one simulation. The dot is the average empirical coverage calculated from 10 simulation repeats. The error bars are $\pm$ 1.96 standard error of mean.

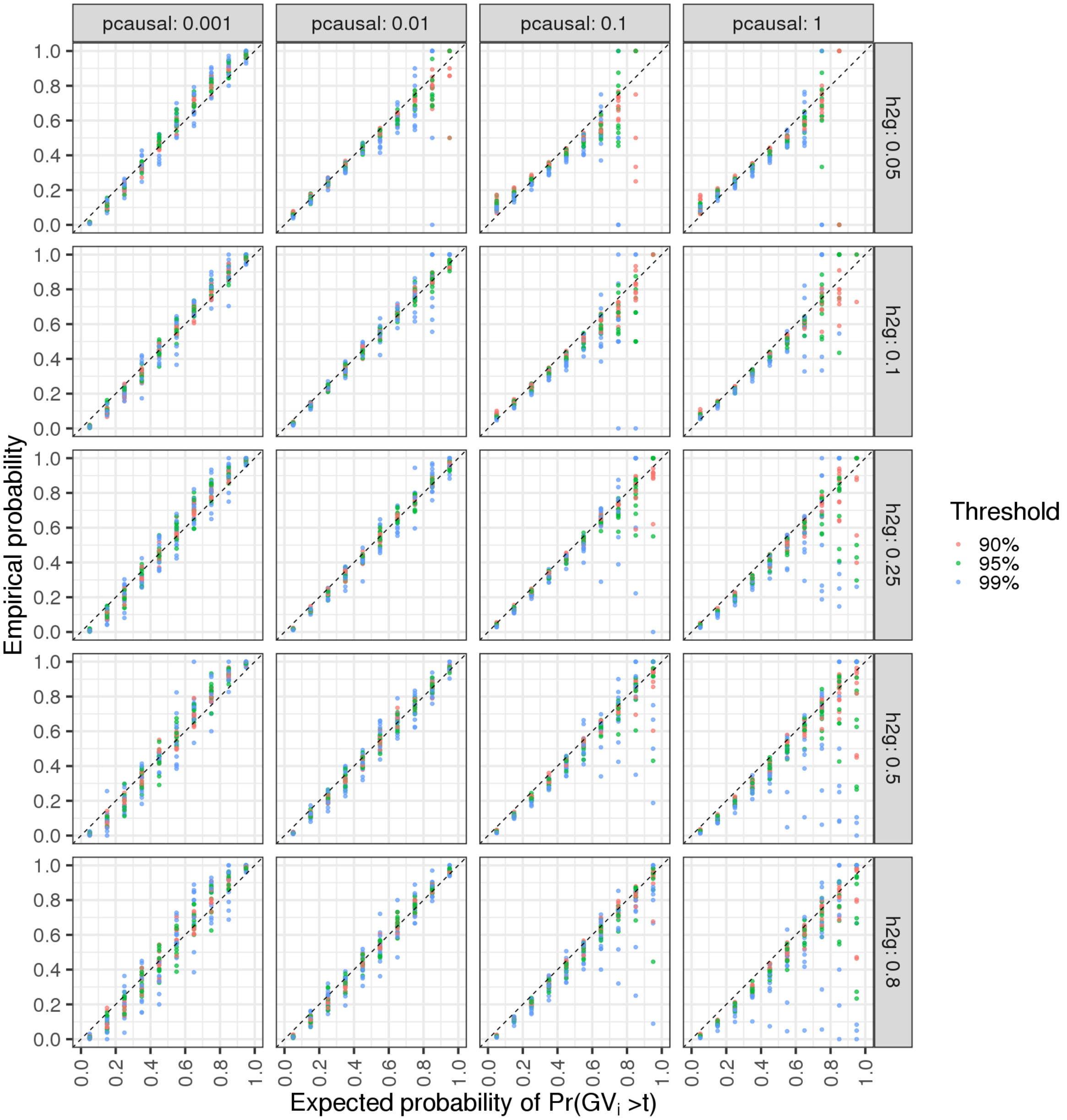


**Supplementary Figure 14. Calibration of probability of GV above threshold t with respect to proportion of causal effects and SNP-heritability**. Individuals are divided into 10 bins (0-1, 0.1 as increment) based on their probability of GV above a prespecified threshold t. The x-axis is expected probability set as middle of each bin. The y-axis is the empirical probability calculated as the proportion of individuals having GV within the lower and upper bound of the bin of one simulation replicate. Each dot represents the expected and empirical probability of corresponding probability bin for one simulation replicate. Different color represents different prespecified threshold.

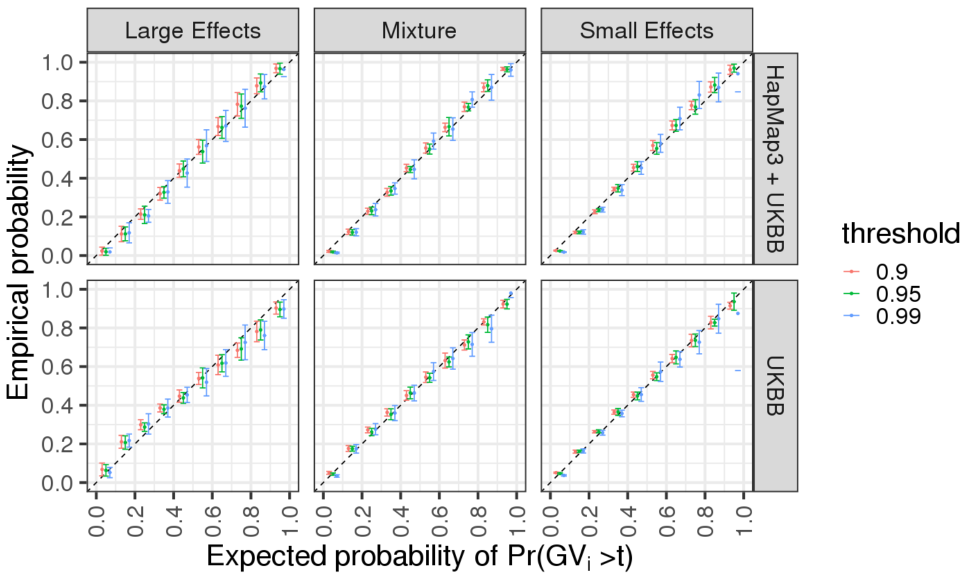


**Supplementary Figure 15. Impact of effect size distribution and SNP panel on calibration of probability of GV above threshold t.** Each column summarizes results for each of the three genetic architectures. Small effects are simulated under $h_g^2 = 0.02, p_{causal} = 0.01$; large effects are simulated under $h_g^2 = 0.02, p_{causal} = 0.001$; Mixture are half and half mixture of the two simulations (small effects: $h_g^2 = 0.01, p_{causal} = 0.005$; large effects: $h_g^2 = 0.01, p_{causal} = 0.0005$). Each row summarizes results of LDpred2 trained with 1) HapMap3 + UKBB SNPs or 2) UKBB array SNPs only. The x-axis is the expected probability set as middle of each bin. The y-axis is the empirical probability calculated as the proportion of individuals having GV within the lower and upper bound of the bin of one simulation replicate. Each dot represents the expected and empirical probability of corresponding probability bin for one simulation replicate. Different colors represent different prespecified thresholds (PRS estimates at each quantile among testing individuals).

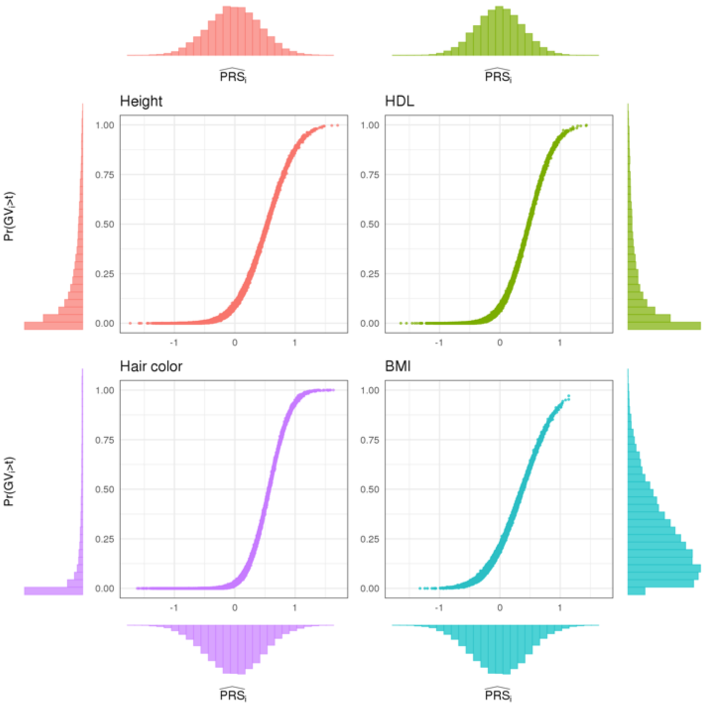


**Supplementary Figure 16. Individual ranking is consistent when ranking by PRS estimates versus probability of genetic value above threshold.** The x-axis is the PRS estimates of testing individuals and the y-axis is the probability that GV is above threshold t, where t is (arbitrarily) set to the 90th percentile in the testing individuals. For the individuals whose PRS estimates are far away from threshold, the probability is 0 and 1 respectively. For individuals close to the stratification threshold, the probability of larger than the threshold increases as PRS estimates increase. The histogram on the x-axis is the distribution of PRS estimates in testing individuals and the histogram on the y-axis is its distribution in testing individuals.

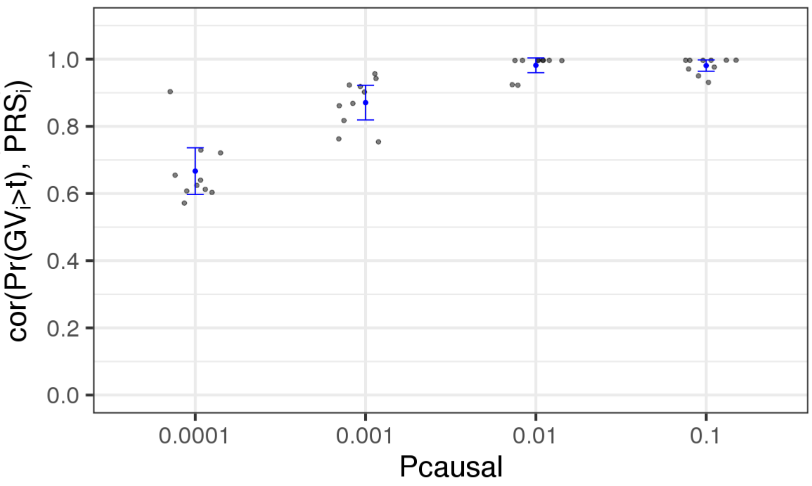


**Supplementary Figure 17. Correlation between PRS and probability of *GV>t* increases as polygenicity increases.** The simulation is performed on 124,080 SNPs on chromosome 2, with heritability equal to 0.2 and polygenicity set to {0.0001, 0.001, 0.01, 0.1}. The grey dots are the Spearman's correlation between PRS and Pr(GV>t) for one simulation replicate. The blue dots and whiskers represent the average Spearman's correlation and standard deviation for each polygenicity.

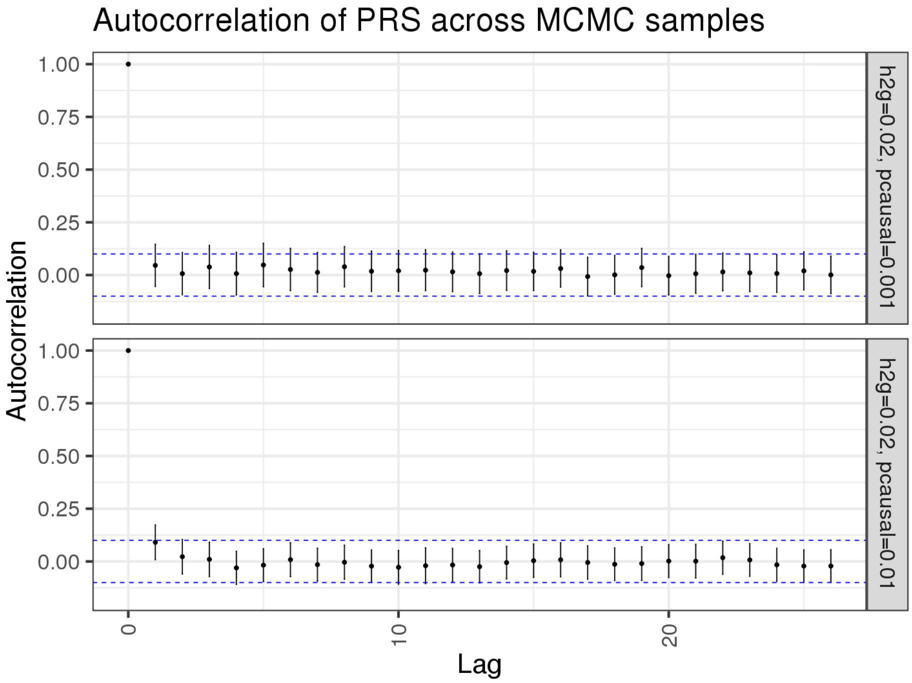


**Supplementary Figure 18. Autocorrelation of PRS samplings across MCMC samples.** For each testing individual, we compute the autocorrelation of the 500 PRS samplings with lag from 0 to 30. The dot is the average autocorrelation at each lag for all 21,723 individuals and the whiskers are 1.96 standard deviations. The simulation is performed on chromosome 2 Hapmap3 + UKBB array SNPs, h2g = 0.02. Upper panel: p_causal = 0.01; lower panel: p_causal = 0.001.
